# Engineering the Immune Adaptor Protein STING as a Biologic

**DOI:** 10.1101/2021.02.18.431824

**Authors:** Xin Sun, Yun Ni, Yanpu He, Mengdi Yang, Tetsuo Tani, Shunsuke Kitajima, David A. Barbie, Jiahe Li

## Abstract

Activation of the stimulator of interferon genes (STING) pathway through cyclic dinucleotides (CDNs) has been explored extensively as potent vaccine adjuvants against infectious diseases as well as to increase tumor immunogenicity towards cancer immunotherapy in solid tumors. Over the last decade, a myriad of synthetic vehicles, including liposomes, polymers, and other nanoparticle platforms, have been developed to improve the bioavailability and therapeutic efficacy of STING agonists in preclinical mouse models. In comparison to synthetic materials, protein-based carriers represent an attractive delivery platform owing to their biocompatibility, amenability to genetic engineering, and intrinsic capacity to form well-defined structures. In the present work, we have engineered the immune adaptor STING as a protein-based delivery system for efficient encapsulation and intracellular delivery of CDNs. Through genetic fusion with a protein transduction domain, the recombinant STING can spontaneously penetrate cells to markedly enhance the delivery of CDNs in a mouse vaccination model and a syngeneic mouse melanoma model. Moreover, motivated by recent findings that certain tumor cells can evade immune surveillance via loss of STING expression, we further unveiled that our STING platform can serve as a functional vehicle to restore the STING signaling in a panel of lung and melanoma cell lines with impaired STING expression. Taken together, our STING-based protein delivery platform may offer a unique direction towards targeting STING-silenced tumors as well as augmenting the efficacy of STING-based vaccine adjuvants.

## INTRODUCTION

The cytosolic DNA sensing pathway involving cyclic GMP-AMP synthase (cGAS) and the stimulator of interferon genes (STING) represents an essential innate immune mechanism in response to foreign pathogens(1). Upon detection of cytosolic DNA, the intracellular nucleic acid sensor cGAS catalyzes the productions of cyclic dinucleotides (CDNs) such as 2’3’-cyclic GMP-AMP (cGAMP), which functions as a second messenger to bind the adaptor protein STING to initiate type I interferon (IFN) production and boost dendritic cell (DC) maturation and T cell infiltration(2). Meanwhile, the cGAS-STING signaling pathway is profound at sensing neoplastic progression by promoting type I IFN production and initiating cytotoxic T cell-mediated anti-tumor immune response(3). These fundamental studies have accelerated the development of utilizing synthetic STING agonists to activate the innate and adaptive immune responses as a monotherapy or in combination with immune checkpoint blockade (ICB) for cancer immunotherapy(4,5).

Despite the promise of CDNs such as cGAMP as immune adjuvants, they suffer from several limitations: (**1**) CDNs exhibit fast clearance from the injection site, which may induce systemic toxicity, (**2**) naturally derived CDNs are susceptible to enzymatic degradation, which can lower the efficacy of adjuvanticity potential, and (**3**) CDNs have inefficient intracellular transport properties due to limited endosomal escape or reliance on the expression of a specific transporter protein(6–8). To address these challenges, existing efforts are largely focused on two main directions: (**1**) generation of novel biomaterial-based delivery systems to improve the *in vivo* delivery of CDNs to activate innate immune cells, and (**2**) discovery of new STING agonist analogs via medicinal chemistry and drug screening to confer greater chemical stability and improved pharmacokinetics(7–11).

Here, we sought to develop a new delivery system that can offer structural simplicity and modularity from the perspective of delivery vehicle design, while becoming an add-on technology by incorporating newly discovered synthetic STING agonist compounds. To this end, we previously uncovered an unnatural function of a recombinant STING protein that lacks the hydrophobic transmembrane (TM) domain (hereinafter referred to as STINGΔTM)(12). Notably, following delivery via commercial transfection reagents, the STINGΔTM/cGAMP complexes can activate the STING signaling pathway even in cells without endogenous STING expression. In our present work, to bypass the need for any synthetic delivery material, we sought to engineer a protein-based carrier for STING agonists by generating a cell-penetrating STINGΔTM (CP-STINGΔTM) through genetic fusion with a cell-penetrating domain, named Omomyc. As a dominant-negative form of the human *MYC* oncogene, Omomyc was originally identified to target *KRAS*-driven tumor cells in several NSCLC xenograft mouse models(13). Intriguingly, in a synthetic vehicle-free mode, CP-STINGΔTM markedly enhanced delivery of cGAMP in cells, which differ in the levels of endogenous STING expression or cell type. To prove its utility *in vivo*, we first explored CP-STINGΔTM to enhance the delivery of cGAMP as an adjuvant in a mouse model vaccinated with chicken ovalbumin(14). Furthermore, in a syngeneic mouse model of melanoma, we explored a combination immunotherapy regimen consisting of an ICB inhibitor, anti-PD-1, and STING agonism(15,16). Collectively, our work demonstrated the potential of repurposing the immune sensing receptor as a vehicle to encapsulate and deliver immune adjuvants towards vaccine and cancer immunotherapy development.

## RESULTS

### Overall Scheme of cGAMP delivery by CP-STINGΔTM

In contrast to existing delivery strategies such as nanoformulations or synthetic depots to overcome the challenges in encapsulation and intracellular delivery of STING agonist (e.g., cGAMP), we have repurposed the natural receptor STING as a highly modular and simple platform to efficiently bind and deliver cGAMP *in vitro* and *in vivo*(7). Specifically, we took advantage of previous biochemical studies, in which the recombinant C-terminal domain of STING protein (STINGΔTM, 139-379aa for human and 138-378aa for mouse) is known to bind cGAMP with high affinity and stability(17,18). Additionally, in our previous work, we serendipitously uncovered that the recombinant STINGΔTM could form complexes with cGAMP, and activate the downstream STING signaling following delivery of the complexes by commercial transfection reagents in HEK293T that do not express endogenous STING. On the contrary, recombinant STINGΔTM proteins with catalytically inactive mutations, including S366A and deletion of the last 9 amino acids (i.e., ΔC9), failed to activate the STING pathway in HEK293T(19,20). Building on this serendipitous discovery, to bypass the need for transfection reagents, here we developed a cell-penetrating (CP)-STINGΔTM to deliver cGAMP into different cell types via genetic fusion of a cell-penetrating protein (**Figure 1a** and **b**). Notably, in contrast to cell-penetrating peptides such as trans-activating transcriptional activator (TAT), we have chosen the Omomyc mini-protein as our cell-penetrating moiety for three reasons: (1) Omomyc (91 amino acids) is derived from a dominant-negative form of the human *MYC* oncogene and has recently shown specific targeting and potent tumor cell penetration capabilities in human cancer cell lines and xenograft mouse models; (2) The natural dimer conformation of Omomyc coincides with STINGΔTM, which also exists as a dimer in the absence of cGAMP; (3) Omomyc may not cause an immunogenicity issue owing to its human origin(13).

**Figure 1.**
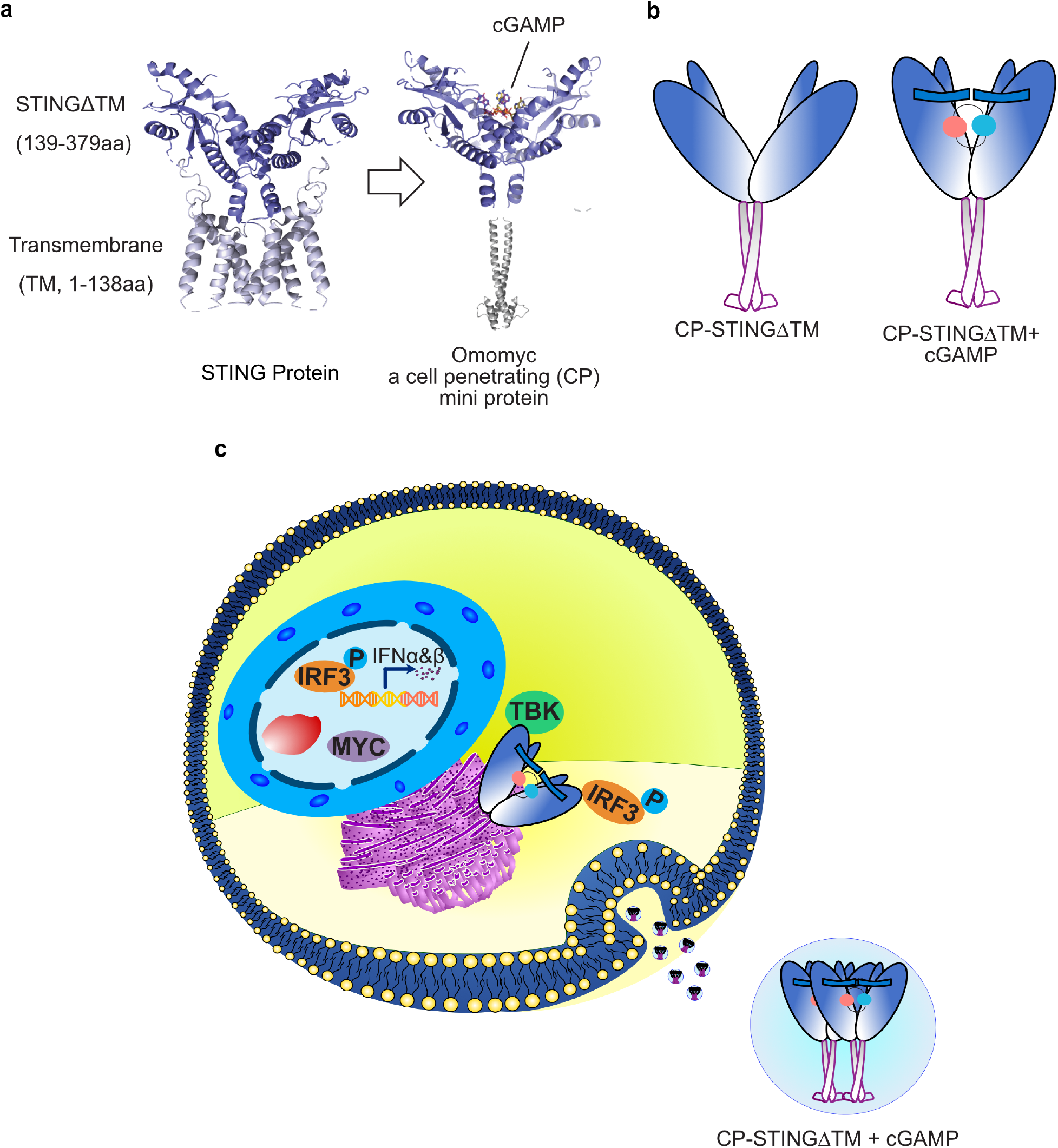
Schematic of using recombinant cell-penetrating (CP)-STINGΔTM as a biologically functional platform for cGAMP delivery. **(a)** To bypass the need for synthetic vehicles, we designed and engineered a CP-STINGΔTM by replacing the transmembrane (TM) of the full-length STING with Omomyc, a cell-penetrating mini protein. **(b)** A cartoon model illustrating how CP-STINGΔTM binds cGAMP. **(c)** By fusing with the cell-penetrating domain, the CP-STINGΔTM is capable of penetrating cells, delivering cGAMP, and engaging with downstream proteins such as TBK1 and IRF3, which result in the production of type I IFNs.

Since the C terminal amino acids of STING directly interact with downstream effector proteins, including TBK1 and IRF3, we genetically fused the cell-penetrating protein Omomyc to the N terminus of STINGΔTM to prevent any steric hindrance posed by Omomyc (**Figure 1c**). In addition, we generated two essential CP-STINGΔTM mutants to help dissect the mechanisms underlying enhanced delivery of cGAMP: one lacks the effector function to engage with the downstream STING signaling pathway and the other fails to bind cGAMP (Table 1)(21–23). After recombinant protein expression in *E. coli*, we purified 6x Histidine (His) tagged proteins via the metal affinity purification and size exclusion chromatography. As shown in **Figure S1**, both size exclusion chromatography studies and SDS-PAGE confirm that the fusion protein can be purified with high yield and homogeneity from *E. coli*. Additionally, the denatured proteins exhibited predicted molecular weights in SDS-PAGE, while the SEC graphs show that CP-STINGΔTM likely forms a tetramer under a native condition in agreement with our previous study(12).

**Table 1:** STING variants used in this study. * Amino acid positions represent the human STING (1-379aa), which are conserved in the mouse STING (1-378aa).

| STING variants* | Description |
| --- | --- |
| STINGΔTM | STING lacking the N terminal transmembrane domain |
| STINGΔTMΔC9 | 9-amino acid deletion at the C terminus that abolishes type 1 IFN induction |
| STINGΔTM(R238A/Y240A) | Deficient for cGAMP binding |
| CP-STINGΔTM | Inclusion of cell-penetrating domain -- Omomyc to bypass transfection reagent |
| CP-STINGΔTMΔC9 | |
| CP-STINGΔTM(R238A/Y240A) | |
| CP-STINGΔTM-dsred | |

### CP-STINGΔTM can effectively internalize cells

While Omomyc protein itself has been shown to internalize different lung cancer cell lines *in vitro* as well as in mouse lung xenografts, it remains to be investigated whether genetic fusion of Omomyc with STINGΔTM can indeed penetrate cells spontaneously. To assess the cell-penetrating potential of CP-STINGΔTM, we treated two human non-small cell lung cancer (NSCLC) cell lines with low or absent STING, H1944 and A549(24), for 24 hours followed by immunostaining against an 8-amino acid FLAG epitope (DYKDDDDK) encoded in between Omomyc and STINGΔTM. Because the FLAG epitope is not known to be expressed by mammalian cells, we could make use of anti-FLAG staining to distinguish exogenously delivered STING protein variants from endogenous STING proteins. Moreover, in contrast to covalently conjugating proteins with fluorescent dyes, which typically modify the surface amine or cysteine groups of proteins, our approach can prevent altering the pharmacokinetics of intracellular protein accumulation. As shown in **Figure 2a** and **c**, CP-STINGΔTM exhibited efficient intracellular uptake in H1944 and A549, while STINGΔTM alone failed to internalize cells owing to the lack of Omomyc to promote cell penetration. In addition, we also genetically fused Omomyc to the catalytically inactive mutant STINGΔTMΔC9, which is known to abolish STING function due to the deletion of 9 amino acids at the very C terminus. As shown in **Figure S2a, c**, and **e**, CP-STINGΔTMΔC9 showed comparable degrees of internalization, which confirmed that the intracellular uptake is mediated by Omomyc instead of STING. To further corroborate our findings beyond fluorescence microscopy, we performed flow cytometry to confirm the uptake profiles of different STING variants after intracellular staining against the same synthetic epitope FLAG (**Figure 2b** and **d**). In addition to the NSCLC cell lines, we validated the uptake of CP-STINGΔTM and CP-STINGΔTMΔC9 in human melanoma and ovarian cancer cell lines by fluorescence microscopy and flow cytometry (**Figure S2b, d,** and **e**). Finally, to dissect the mechanism by which the cell-penetrating STINGΔTM enters cells, we tested a range of small molecule inhibitors targeting different endocytic pathways including: 5-(N-Ethyl-N-isopropyl) amiloride (EIPA), chlorpromazine, Dynasore, cyclodextrin, and Filipin. Among the small molecule inhibitors we have tested, a macropinocytosis inhibitor, EIPA and an endocytosis inhibitor, Dynasore exhibited a dose-dependent inhibition of CP-STINGΔTM in H1944 (**Figure 2e-g** and **Figure S2f,g**)(13,25). In contrast, inhibitors targeting other uptake pathways failed to inhibit the uptake of CP-STINGΔTM (**Figure S2h**, and **i**). Of note, our findings agree with the previous work, in which the Omomyc protein itself was taken up by cancer cells primarily through the macropinocytosis and endocytosis pathways(13). Therefore, we conclude that the cell-penetrating capability of the fusion protein is mediated by Omomyc in a macropinocytosis and endocytosis-dependent manner.

**Figure 2.**
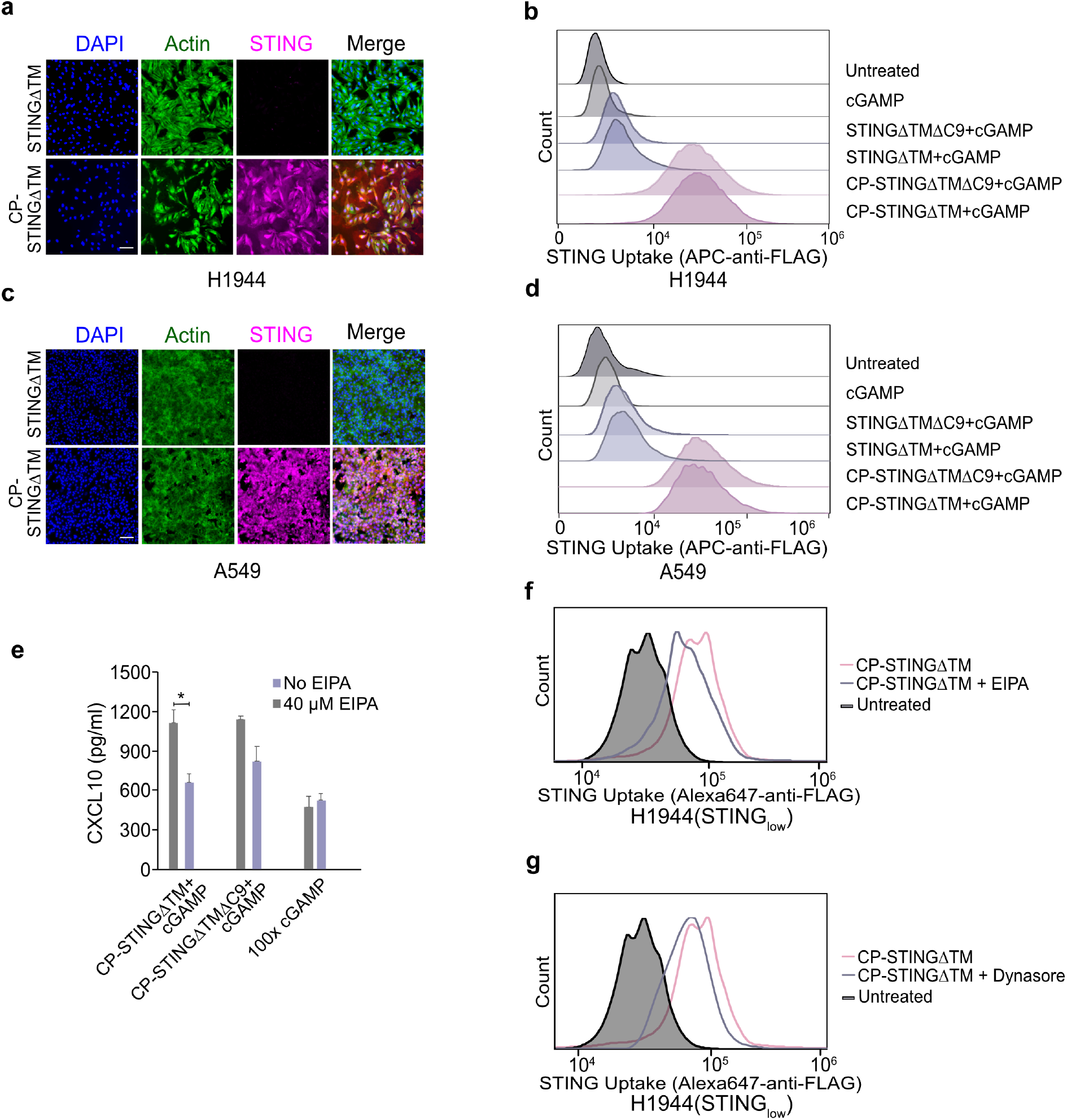
CP-STINGΔTM spontaneously internalizes cancer cells. Fluorescence microscopy imaging of internalized CP-STINGΔTM in H1944 (STING_low_) with downregulated STING expression **(a)** and A549 (STING_absent_) without any STING expression **(c)** (scale bar = 100 μm). Flow cytometry of internalized CP-STINGΔTM in H1944 (STINGlow) with downregulated STING expression **(b)** and A549 (STING_absent_) without any STING expression (d). Cells were treated with “40 μg/ml CP-STINGΔTM” + 1 μg/ml cGAMP” or “40 μg/ml STINGΔTM + 1 μg/ml cGAMP” for 24 hours before staining with APC-anti-FLAG. **(e)** H1944(STING_low_) were preincubated with 40μM EIPA for 2 hours and treated with “40 μg/ml CP-STINGΔTM + 1 μg/ml cGAMP” or “40 μg/ml STINGΔTM + 1 μg/ml cGAMP” or 100 μg/ml cGAMP, CXCL10 production was inhibited by macropinocytosis inhibitor EIPA. Representative flow cytometry analysis of CP-STINGΔTM uptake in H1944 pretreated with indicated inhibitors targeting macropinocytosis (f) and endocytosis (g), respectively.

### CP-STINGΔTM enhances cGAMP delivery and STING activation in a panel of lung and melanoma cell lines with impaired STING expression

In contrast to innate immune cells, which are highly sensitive to cGAMP-mediated STING activation, previous work by others have shown that downregulation of STING in tumor cells greatly reduced the sensitivity of cancer cells to STING agonists, which can promote immune suppression and exclusion of cytotoxic T cells in the tumor microenvironment (TME)(24,26). Therefore, we sought to ask whether the fusion protein could promote intracellular delivery of the STING agonist cGAMP in a panel of cell lines with reduced sensitivity to STING agonists. We first focused on two STING_low_ NSCLC cell lines, H1944 and H2122, in which the expression of endogenous STING is downregulated due to histone methylation at the native STING promoter(24). As shown in **Figure 3a** and **S3c**, we compared “CP-STINGΔTM + cGAMP”, “CP-STINGΔTMΔC9 + cGAMP”, free cGAMP, and lipofectamine-transfected cGAMP to vehicle control-treated cells. Of note, a 1:1 molar ratio of one STING dimer to one cGAMP was prepared for different STING/cGAMP complexes. Impressively, the codelivery systems comprising “CP-STINGΔTM + cGAMP” or “CP-STINGΔTMΔC9 + cGAMP” required ~100-fold lower concentration of cGAMP than free cGAMP or lipofectamine-transfected cGAMP to induce comparable levels of CXCL10, one of the chemokines that can be induced by the STING pathway(27,28). In addition, since the STING activation in tumor cells can upregulate major histocompatibility complex I (MHC-I) to promote cytotoxic T cell recognition, we measured the surface expression of MHC-I in the same cancer cells(29). Consistent with measurement of CXCL10 by ELISA, “CP-STINGΔTM + cGAMP” and “CP-STINGΔTMΔC9 + cGAMP” similarly enhanced surface expression of MHC class I in H1944 and melanoma cells (**Figure S3d** and **e**).

**Figure 3.**
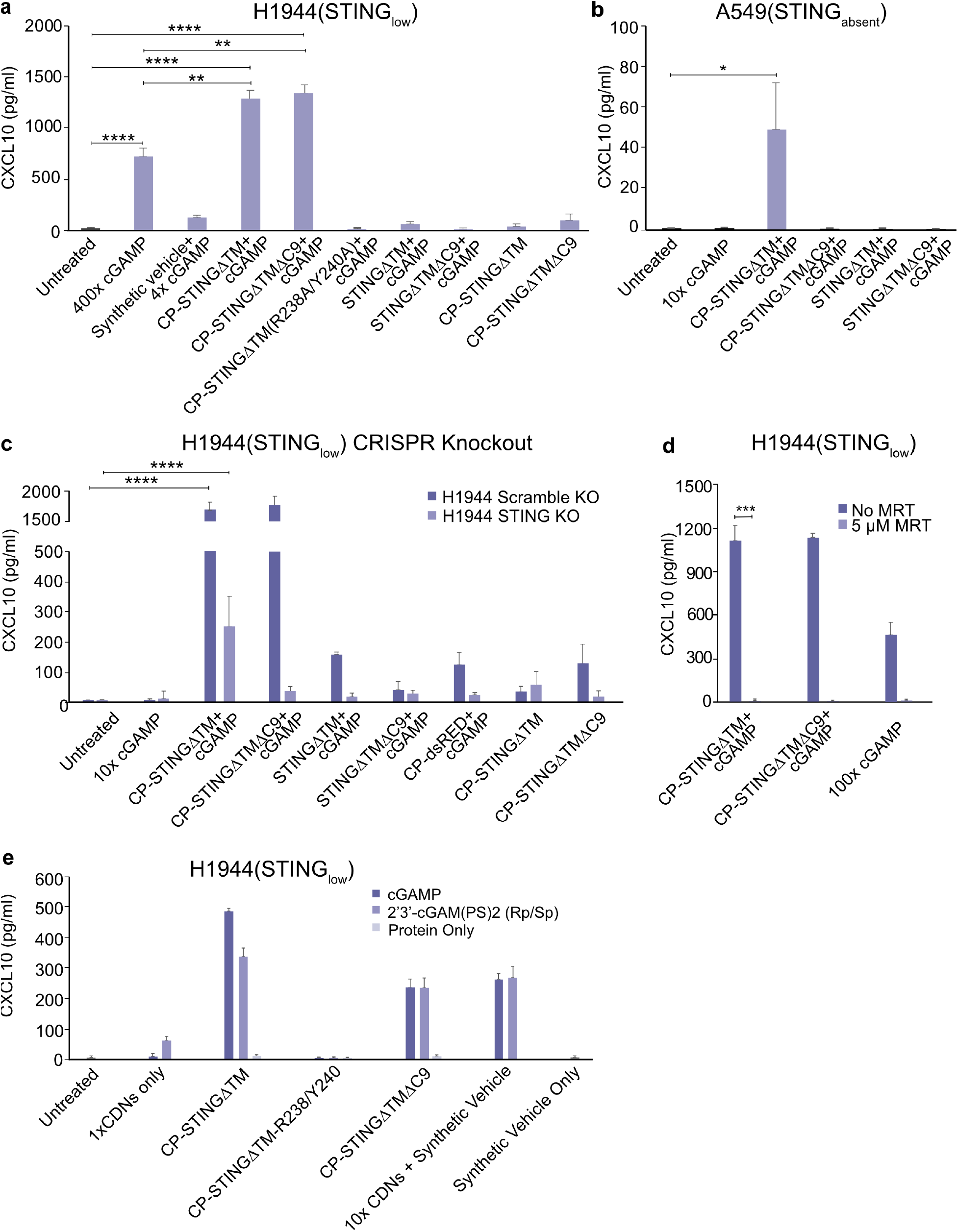
CP-STINGΔTM markedly enhances cGAMP delivery and STING activation in cancer cell lines with impaired STING expression. (**a**) CP-STINGΔTM plays a chaperon role in H1994 (STING_low_) that have down-regulated STING expression. Specifically, CXCL10 was remarkably enhanced by “10 μg/ml CP-STINGΔTM + 0.25 μg/ml cGAMP” or “10 μg/ml CP-STINGΔTMΔC9 (catalytically inactive mutant) + 0.25 μg/ml cGAMP” compared to 100-400 fold higher concentration of free cGAMP and 40 fold higher concentration of cGAMP delivered by Lipofectamine 2000. (**b**) “CP-STINGΔTM + cGAMP” forms a functional complex in A549 (STING_absent_), which does not express endogenous STING. Only “40 μg/ml CP-STINGΔTM +1 μg/ml cGAMP” could induce CXCL10. (**c**) After knocking out endogenous STING in H1944 by CRISPR, CXCL10 expression was only induced by “40 μg/ml CP-STINGΔTM + 1 μg/ml cGAMP” but not by the catalytic inactive “40 μg/ml CP-STINGΔTMΔC9 + 1 μg/ml cGAMP” or free cGAMP. (**d**) The CXCL10 production was inhibited by the TBK1 inhibitor - MRT, which indicates that the enhanced STING signaling by CP-STINGΔTM or CP-STINGΔTMΔC9 was dependent on the TBK1, a key component in the STING pathway. (**e**) Codelivery of CP-STINGΔTM and a synthetic, non-degradable cGAMP analog, cGAMP(PS)_2_(Rp/Sp), also enhances CXCL10 production in comparison to free cGAMP(PS)_2_(Rp/Sp) or 10x cGAMP(PS)_2_(Rp/Sp) transfected by Lipofectamine 2000, which suggests that CP-STINGΔTM promotes the cGAMP delivery instead of protecting cGAMP from enzymatic degradation. *P<0.05, **P<0.01, ***P<0.001, ****P<0.0001. Values = mean ± SEM, n=4.

To explain our findings, we first ruled out the possibility of endotoxin contamination resulting from protein purification from *E. coli*, as CP-STINGΔTM or CP-STINGΔTMΔC9 protein alone of equivalent concentrations did not induce CXCL10 (**Figure 3a**). It is intriguing, however, delivery of cGAMP by the catalytically inactive CP-STINGΔTMΔC9, in which the interaction of STING with TBK1 and IRF3 is disabled, enhanced the STING activation to a degree similar to that of the wildtype (i.e., CP-STINGΔTM) (**Figure 3a**). We hypothesized that in the STING_low_ cell lines H1944 and H2122, the CP-STINGΔTM primarily may serve as a chaperon by promoting delivery of cGAMP into tumor cells. To test this hypothesis, we generated two additional fusion proteins: CP-dsRed and CP-STINGΔTM (R238A/R240A). Importantly mutations of the 238th arginine (R238) and 240th tyrosine (Y240) to alanine (A) are known to abolish the ability of STING to bind cGAMP(23). As shown in **Figure 3a**, **S3d**, and **e**, these two protein variants failed to enhance CXCL10 production to the same extent as “CP-STINGΔTM + cGAMP” and “CP-STINGΔTMΔC9 + cGAMP”. Therefore, through genetic mutations that inactivate two separate functions of STING, including the effector and cGAMP-binding capabilities, we have found that in STING_low_ cells, CP-STINGΔTM primarily act as a chaperon to efficiently deliver cGAMP intracellularly and therefore greatly enhancing the STING activation.

Motivated by the ability of CP-STINGΔTM to markedly enhance cGAMP delivery and STING activation in STING_low_ cells, we further extended our observations to A549 (human NSCLC) and SK-MEL-5 (human melanoma), which do not express endogenous STING (STING_absent_)(24,30). Interestingly, we found that only “CP-STINGΔTM + cGAMP” induced CXCL10, while the catalytically inactive CP-STINGΔTMΔC9 along with cGAMP did not (**Figure 3b**). Additionally, “STINGΔTM + cGAMP” failed to induce CXCL10, which can be explained by the absence of Omomyc to facilitate cell penetration (**Figure 3a**). These observations imply that codelivery of CP-STINGΔTM and cGAMP functionally restored the deficient STING signaling in STING_absent_ cells. To further confirm this hypothesis, we utilized Clustered Regularly Interspaced Short Palindromic Repeats (CRISPR) to genetically knock out endogenous cGAS and STING, respectively in H1944. Notably, the cGAS knockout is known to inhibit the production of endogenous cGAMP(31). Consistent with data in STING_low_ cell lines, in H1944 with cGAS knockout but intact STING, both “CP-STINGΔTM + cGAMP” and “CP-STINGΔTMΔC9 + cGAMP” could comparably induce CXCL10 expression, suggesting that endogenous cGAMP is not required for the activation of STING signaling (**Figure S3f**). In H1944 with only STING knockout, however, CXCL10 expression was induced by “CP-STINGΔTM + cGAMP” but not the catalytically inactive “CP-STINGΔTMΔC9 + cGAMP” (**Figure 3c**), which is consistent with findings in A549 and SK-MEL-5 cells, in which endogenous STING expression is completely absent (**Figure 3b** and **S3f**). In addition, concurrent treatment with a TBK1 inhibitor, MRT, repressed the production of CXCL10 in the cells treated with “CP-STINGΔTM + cGAMP” and “CP-STINGΔTMΔC9 + cGAMP” (**Figure 3d**)(12). Therefore, through both genetic and pharmacological inhibition of key proteins in the STING pathway, we have shown that “CP-STINGΔTM + cGAMP” acts as a functional complex to induce STING signaling in the cells lacking endogenous STING expression. Finally, since cGAMP can be degraded by Ectonucleotide pyrophosphatase/phosphodiesterase 1 (ENPP1), which is abundant in extracellular and intracellular environments, another possibility for enhanced cGAMP delivery is that CP-STINGΔTM may protect cGAMP from ENPP1-mediated hydrolysis(8). To test this possibility, we explored cGAM(PS)_2_(Rp/Sp), a synthetic non-degradable cGAMP analog, in H1944, and observed that “CP-STINGΔTM + cGAM(PS)_2_(Rp/Sp)” and “CP-STINGΔTMΔC9 + cGAM(PS)_2_(Rp/Sp)”, markedly enhanced CXCL10 production in comparison to cGAM(PS)_2_(Rp/Sp) alone of equivalent concentration or at a 10x concentration transfected by a commercial transfection reagent. Moreover, CP-STINGΔTM (R238A/R240A), in which the two mutations R238A and R240A abolish the cGAMP binding, failed to enhance CXCL10 production in the codelivery with cGAM(PS)_2_(Rp/Sp) (**Figure 3e**).

### CP-STINGΔTM improves the efficacy of cGAMP as an immune adjuvant

cGAMP has been explored as a potent vaccine adjuvant that promotes both humoral and cellular immune responses in different mouse vaccination models(32). However, free cGAMP is prone to fast clearance and degradation owing to low molecular weight (~600 Da) and the presence of hydrolyzable phosphoester bonds, respectively. To address these limitations, a myriad of synthetic biomaterials have been developed to enhance the delivery efficacy of cGAMP. In our own work, motivated by enhanced activation of the STING pathway by CP-STINGΔTM in different cell types, we ask whether it could serve as a protein-based delivery platform to efficiently deliver cGAMP as an immune adjuvant. To this end, we made use of the murine dendritic cell line DC 2.4 as a model of antigenpresenting cells (APCs)(7). Similar to our findings in cancer cells, it was shown that CP-STINGΔTM + cGAMP greatly induced expression of CXCL10 and surface expression of MHC-I compared to free cGAMP (**Figure 4a** and **b**).

**Figure 4.**
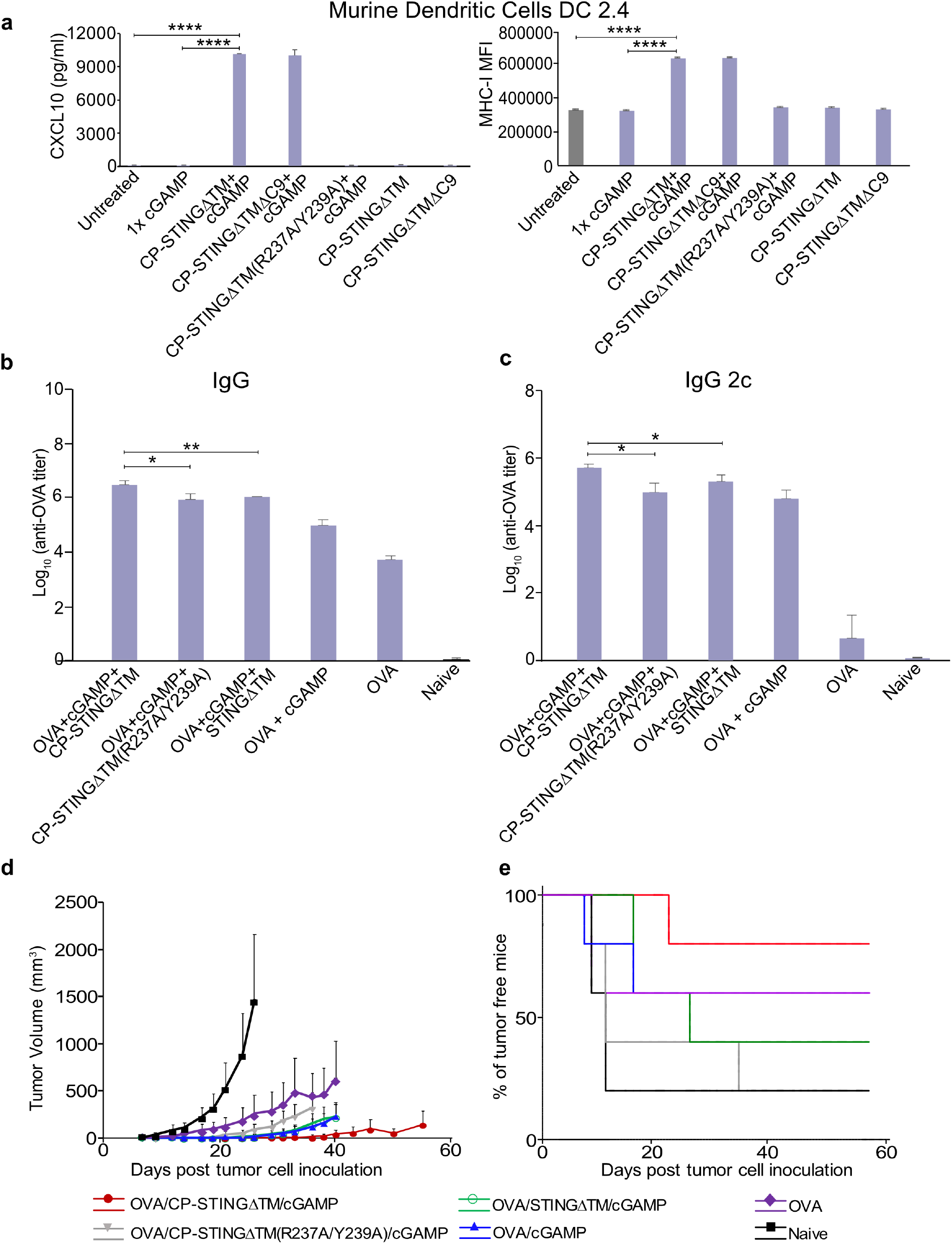
CP-STINGΔTM enhances the efficacy of cGAMP as an adjuvant. (**a**) In murine dendritic cells DC 2.4, “40 μg/ml CP-STINGΔTM + 1 μg/ml cGAMP” markedly induced CXCL10 expression as evidenced by ELISA as well as upregulated surface expression of MHC-I measured by flow cytometry. Levels of OVA-specific total IgG (**b**) and the type I IFN-associated subtype IgG2c (**c**) in groups of C57BL/6 mice (n=5). Mice were immunized with OVA alone, or OVA mixed with 1 μg/ml free cGAMP or combinations of 40 μg/ml STINGΔTM variants with or without 1 μg/ml cGAMP on days 0 and 14 via tail-based injection. On days 21, sera from different vaccination combinations were collected for OVA-specific total IgG and IgG2c quantification. On day 21, the same cohort of mice were challenged with 1 million B16-OVA (257-264aa) subcutaneously. Data of overall tumor growth (**d**), with survival rate (**e**) at the end of the study were denoted. Values are reported as mean ± SEM. Statistical analysis was performed by one-way ANOVA according to the scales of *P<0.05, **P<0.01, ***P<0.001, and ****P<0.0001.

Next, we tested our hypothesis in wild-type C57BL/6 mice by vaccinating them with a model antigen, chicken ovalbumin (OVA), along with free cGAMP or cGAMP + CP-STINGΔTM serving as an immune adjuvant(12). Following a priming-boost protocol with a two-week interval, we quantified the levels of OVA-specific total IgG as well as type I IFN-associated IgG2c from mouse serum, of which the latter IgG subtype can be induced through activation of the STING pathway. As shown by the OVA-specific ELISA, the “OVA + cGAMP + CP-STINGΔTM” treatment group increased the levels of OVA-specific IgG and IgG2c by nearly ten-fold compared to “OVA + cGAMP + CP-STINGΔTM (R237A/Y239A)”, “OVA + cGAMP + STINGΔTM”, and “OVA + cGAMP” (**Figure 4b and c, S4a-d**). To examine the cellular responses, we measured the percentage of CD8 T cells carrying the MHC-I-SIINFEKL epitope (OVA_257-264aa_) via tetramer staining (**Figure S4b**). In agreement with studies in humoral responses, “OVA + cGAMP + CP-STINGΔTM” exhibited the highest induction of SIINFEKL-specific CD8 T cells among different treatment groups. Furthermore, when comparing CP-STINGΔTM to STINGΔTM, the latter of which does not have the cell-penetrating protein domain, CP-STINGΔTM markedly enhanced OVA-specific IgG and IgG2c as well as SIINFEKL-restricted CD8 T cells (**Figure 4b, c**, and **S4a-d**)(33). We reasoned that it is due to increased retention and intracellular uptake mediated by the cell-penetrating protein Omomyc. Indeed, in a later experiment, we found that CP-STINGΔTM exhibited greater retention in tumors than STINGΔTM at 96 hours post injection (**Figure 6a** and **b**). Next, we made use of the same cohort of vaccinated C57BL/6 mice to examine whether the increased induction in antigen-specific IgG and CD8 levels could confer greater protection in a prophylactic syngeneic mouse melanoma model. Specifically, one week after the boost, we challenged the mice with B16 melanoma cells engineered to express the SIINFEKL epitope. As shown in **Figure 4d** and **e**, the cohort vaccinated with “OVA + cGAMP + CP-STINGΔTM” combination displayed the slowest tumor growth rates and longest survival rates.

**Figure 5.**
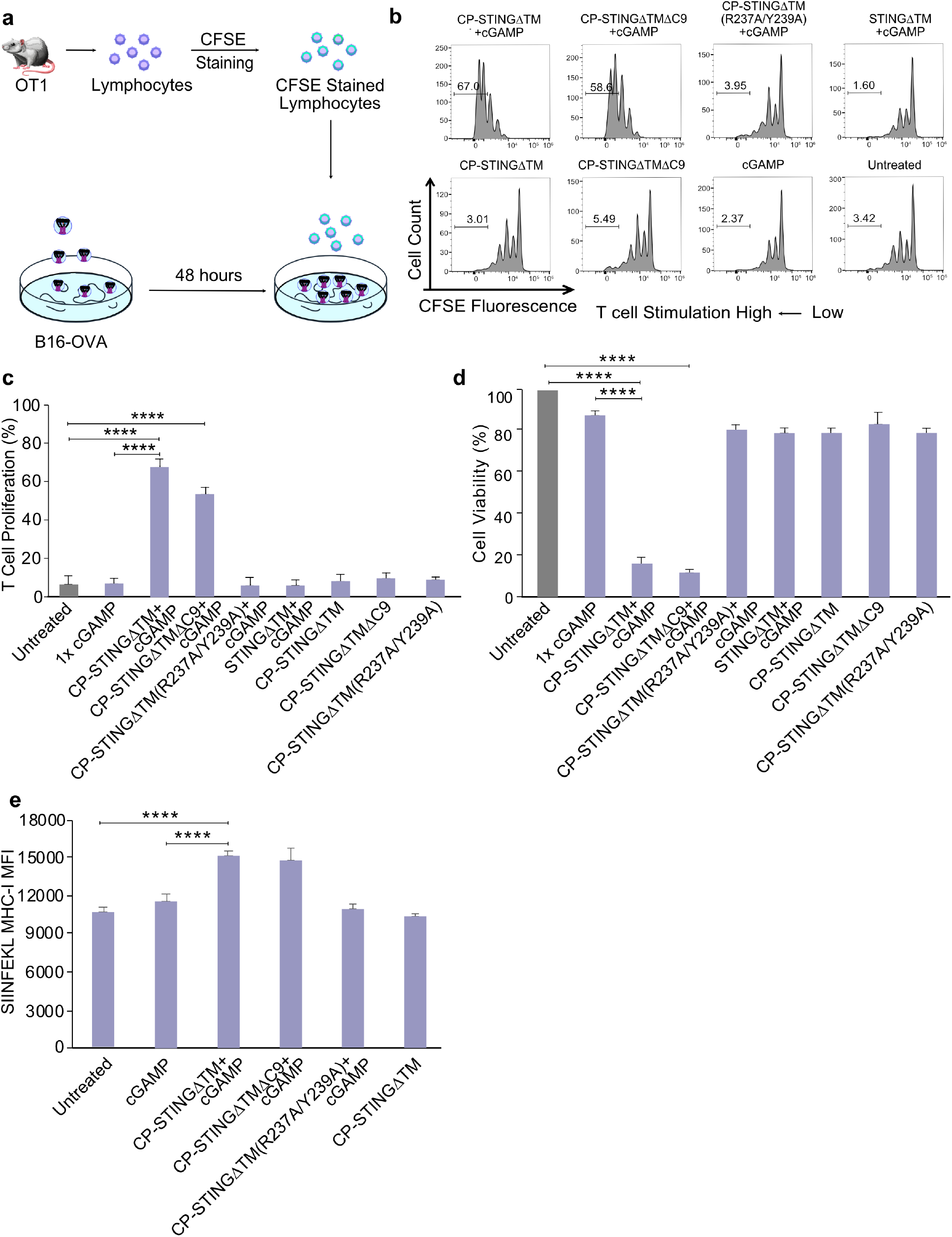
*Ex vivo* T cell-mediated cancer cell killing after activating the STING pathway in tumor cells. **(a)** CFSE-labeled OT1 cells were added into B16-OVA (257- 264aa) cells that were pretreated with cGAMP plus indicated STINGΔTM variants for 48 hours (~10:1 ratio of effector T cell to tumor cells). Proliferated T cells were assayed five days later. **(b)** Representative CFSE flow cytometry data from one of four independent experiments are displayed. **(c)** Quantification of T cell proliferation by CFSE staining. While the pretreatment groups “40 μg/ml CP-STINGΔTM + 1 μg/ml cGAMP” and “40 μg/ml CP-STINGΔTMΔC9 + 1 μg/ml cGAMP” promoted T cell proliferation, the variants with deficiency in cGAMP binding or cell penetration did not. **(d)** OT1-mediated cancer cell killing. B16-OVA (257-264aa) that had been pretreated with indicated STING variants plus cGAMP for 48 hours, were cocultured with OT1 cells. After five days, nonadherence T cells were removed by washing, and the viability of adherent tumor cells was assessed by the MTT assay. Experiments were repeated three times. **(e)** Upregulation of SIINFEKL-restricted MHC-I on the surface of B16-OVA (257-264aa). After treating tumor cells with 1 μg/ml cGAMP and 40 μg/ml STING variants for 48 hours, only “40 μg/ml CP-STINGΔTM + 1 μg/ml” cGAMP and “40 μg/ml CP-STINGΔTMΔC9 + 1μg/ml cGAMP” upregulated the expression of SIINFEKL-restricted MHC-I. Graphs are expressed as mean ± SEM (n=4) and statistical analysis by one-way ANOVA according to the following scale: *P<0.05, **P<0.01, ***P<0.001, and ***P<0.0001.

**Figure 6.**
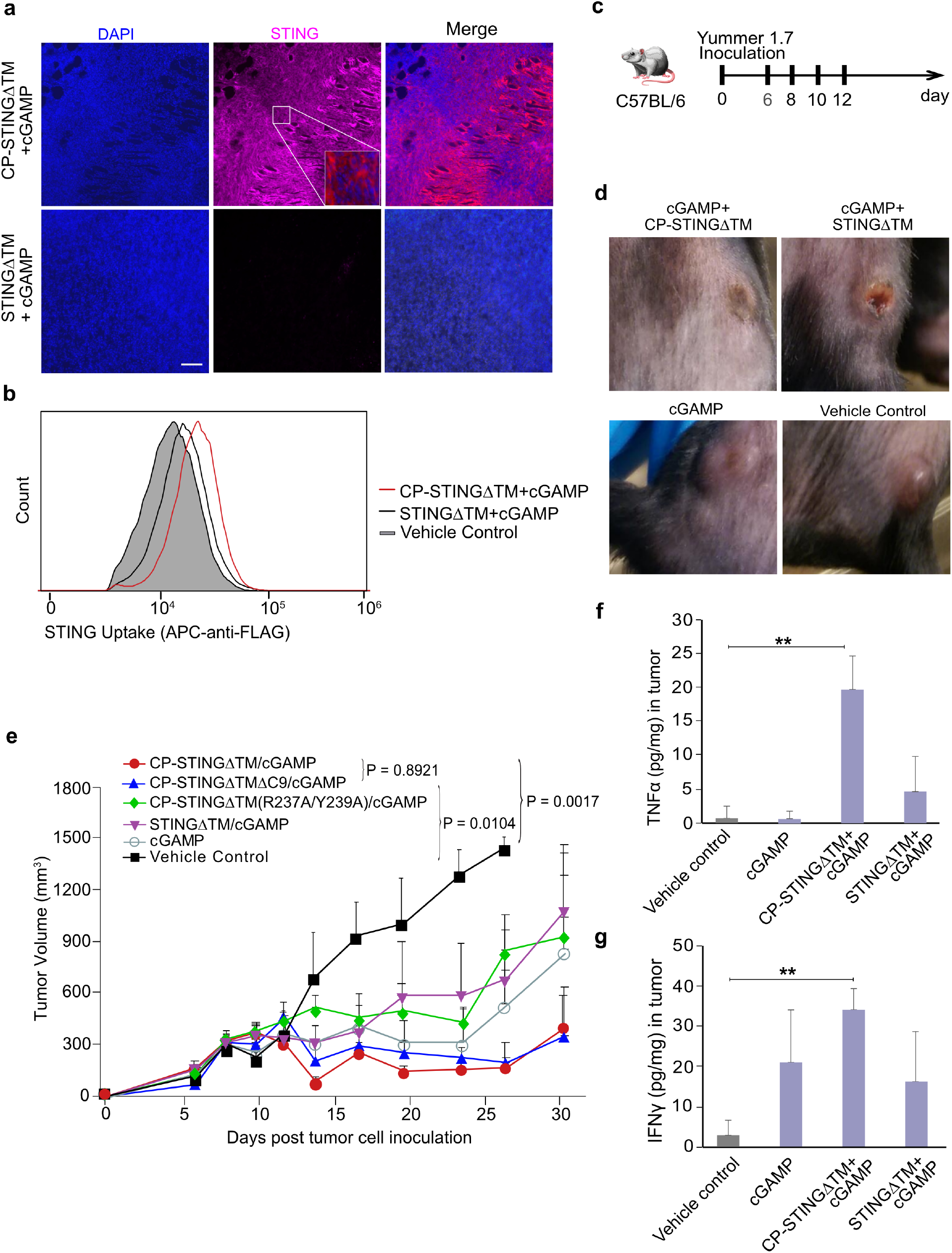
Combining CP-STINGΔTM/cGAMP and anti-PD-1 in a syngeneic mouse melanoma model. Groups of C57BL/6 mice were inoculated with 1 million YUMMER1.7 melanoma cells in the flank and when tumors reached ~150 mm^3^, mice were treated with intraperitoneal injection of anti-PD-1 (200 μg per mouse) and concurrently with intratumoral injection of “100 μg/ml CP-STINGΔTM + 2.5 μg/ml cGAMP” (n = 5), “100 μg/ml CP-STINGΔTMΔC9 + 2.5 μg/ml cGAMP” (n=5), “100 μg/ml CP-STINGΔTM(R237A/Y239A) +cGAMP” (n=5), “2.5 μg/ml cGAMP only” (n=5), and vehicle control (n= 4) (**c**). Cellular uptake of CP-STINGΔTM (n=2) was evaluated with (**a**) microscopic imaging and (**b**) flow cytometry. (**d**) Photos for acute responses for the treatment were taken 72 hours after treatment. (**e**) Overall tumor growth curves were measured using clipper, and tumor volume was calculated using formulations V = (L x W x W)/2, where V is tumor volume, L is tumor length, and W is tumor width. Expression of TNF-alpha (**f**) and IFN-gamma (**g**) induced by various treatment groups (n=3) was quantified by ELISA. Statistical analysis was performed by one-way ANOVA: *P<0.05, **P<0.01.

### Codelivery of CP-STINGΔTM and cGAMP augments tumor cell killing by antigenspecific T cells *ex vivo*

In addition to promoting maturation and cross-presentation of dendritic cells for T cell priming, which serves as the very first step of immune clearance of tumor cells, activation of the STING pathway in tumor cells has been shown to augment cytotoxic T cell-mediated cancer cell killing by upregulating MHC-I on the surface of tumor cells(29). Motivated by the aforementioned vaccination and prophylactic mouse model studies, we next explored whether CP-STINGΔTM and cGAMP can enhance tumor cell killing. To this end, in an *ex vivo* model, we generated two isogenic B16 melanoma cell lines expressing either SIINFEKL-GFP fusion or GFP alone, and treated them with free cGAMP, “cGAMP + CP-STINGΔTM”, “cGAMP + CP-STINGΔTMΔC9” and “cGAMP + CP-STINGΔTM (R237Y239A)” for 48 hours(34). After the supernatant was removed from the tumor cells, CFSE-stained SIINFEKL-specific CD8 T cells, which were harvested from lymph nodes of OT-1 mice, were cocultured with tumor cells (**Figure 5a**). It is noteworthy that by pretreating tumor cells with cGAMP and different STING protein variants followed by washing and coculturing with antigen-specific T cells, we specifically tested the effects of STING activation in tumor cells. As shown in **Figure 5b** and **c**, following a 120-hour coculture, cGAMP complexed with CP-STINGΔTM and CP-STINGΔTMΔC9 induced the highest T cell proliferation as evidenced by T cell division-mediated CFSE dilution in flow cytometry. Moreover, the highest efficacy of tumor killing was detected in the same treatment groups by staining viable tumor cells with MTT after washing away nonadherent T cells (**Figure 5d**). Of note, the tumor killing was only detectable in B16 cells bearing the SIINFEKL epitope but not in the GFP-expressing B16 cells in the coculture with OT-1 cells, indicating that the increased T cell proliferation and tumor cell killing were antigenspecific (**Figure S5a** and **b**). To confirm that the increased T cell proliferation and killing resulted from enhanced tumor cell recognition by OT-1 T cells, after treating SIINFEKL-expressing B16 with cGAMP and different STING variants for 48 hours, we quantified the expression levels of MHC-I and SIINFEKL-restricted MHC-I on the surface of tumor cells by flow cytometry. As shown in **Figure 5e** and **S5c**, only “CP-STINGΔTM + cGAMP” and “CP-STINGΔTMΔC9 + cGAMP” markedly upregulated the expression of MHC-I and SIINFEKL-restricted MHC-I in comparison to free cGAMP and other control treatment groups. We reason that since B16 cells express endogenous STING (**Figure S3a**), CP-STINGΔTM acted as a chaperon to enhance cGAMP delivery into tumor cells in this setting.

### Codelivery of CP-STINGΔTM and cGAMP enhances the therapeutic efficacy of ICB

Having validated the enhanced tumor cell killing by OT-1 cells via codelivery of CP-STINGΔTM and cGAMP *ex vivo*, we further examined whether this approach could augment the efficacy of the combination immunotherapy involving STING agonism and ICB. Here, we made use of an immunogenic mouse melanoma cancer model bearing YUMMER1.7 tumor cells for three reasons: First, YUMMER1.7 cells carry *Braf* mutation and *Pten* loss that mimic the most frequent mutations happening in melanoma patients(35). Second, tumors with increased immunogenicities are generally responsive to ICB, such as anti-PD-(L)1, among which lung cancer and melanoma are of high mutation burden(36). Third, STING activation in the TME has been shown to improve the therapeutic efficacy of ICB in different syngeneic mouse cancer models(37).

Before the treatment study, we first confirmed that CP-STINGΔTM can internalize tumor cells and other cell types in the TME. Specifically, when YUMMER1.7 tumors reached ~150 mm^3^ in C57BL/6 mice, a single dose of CP-STINGΔTM was administered intratumorally. Mice were sacrificed at 96 hours, and tumors were harvested for cryosectioning and immunostaining using the anti-FLAG antibody specific for recombinant STING protein variants. As shown in **Figure 6a**, CP-STINGΔTM but not STINGΔTM was readily detectable across different tumor slices in a homogeneous pattern at 96 hours after a single intratumoral administration, and CP-STINGΔTM was primarily localized in the cytoplasm, suggesting that the presence of the cell-penetrating domain Omomyc domain facilitated the retention of recombinant STING in the TME. To corroborate this finding, in a separate cohort of mice, single cells were prepared for intracellular staining against the same FLAG epitope. Similar to our *in vitro* cellular uptake studies, CP-STINGΔTM efficiently internalized tumor cells in comparison to STINGΔTM that lacks the cell-penetrating capability (**Figure 6b**).

Next, we investigated the therapeutic efficacy of CP-STINGΔTM and cGAMP in combination with anti-PD1 in the Yummer1.7 syngeneic mouse model (**Figure 6c**). Of note, we initiated treatment in mice with relatively large subcutaneous tumors, which are more challenging to treat with immunotherapy than smaller tumors(7). After tumors reached ~150mm^3^, CP-STINGΔTM, CP-STINGΔTMΔC9, CP-STINGΔTM(R237A/Y239A) and STINGΔTM were intratumorally administered with cGAMP, while anti-PD1 was given intraperitoneally at optimized doses every two days for a total of four treatments (**Figure 6c**). Over the duration of treatment, no significant weight loss was detected among different treatment groups in comparison to the vehicle control group (**Figure S6a**). Importantly, both CP-STINGΔTM and CP-STINGΔTMΔC9 showed marked reduction in the tumor progression compared to CP-STINGΔTM(R237A/Y239A) and STINGΔTM treatment groups (**Figure 6d** and **e**). These findings agree with our studies *in vitro*: (1) The mutations R237A/Y239A in STING abolish the binding of cGAMP, and therefore CP-STINGΔTM(R237A/Y239A) cannot effectively deliver cGAMP into target cells. (2) STINGΔTM alone cannot efficiently penetrate target cells due to the absence of the Omomyc protein. (3) because cancer cells and hematopoietic cells in tumors express endogenous STING, CP-STINGΔTM plays a chaperon role in enhancing the intracellular delivery of cGAMP such that there was no detectable difference between CP-STINGΔTM and CP-STINGΔTMΔC9, the latter of which cannot activate the STING signaling. In addition to tumor volume therapeutic efficacy, we further measured proinflammatory cytokines in a separate cohort of mice bearing the same tumor cells(38). The treatment group of “CP-STINGΔTM + cGAMP” displayed increased expression of TNFα, IFNγ, and CXCL10 in comparison to “STINGΔTM + cGAMP” and the untreated group (**Figure 6f, g** and **Figure S6b**).

## DISCUSSION

In this study, we have successfully developed a protein carrier (CP-STINGΔTM) for efficient cytosolic delivery of STING agonists by merging the inherent capacity of the transmembrane deleted STING (STINGΔTM) in binding cGAMP and activating the downstream STING signaling with the cell-penetrating (CP) miniprotein Omomyc, the latter of which was recently validated in several preclinical NSCLC mouse xenograft model as an anti-cancer therapy. Importantly, while the N terminus of Omomyc is responsible for cell targeting, the C terminus of STINGΔTM is involved in intracellular STING functions(19). Additionally, the two protein domains exist as a dimer on their own. Therefore, the fusion protein consisting of CP and STINGΔTM can in theory function properly with the natural configuration and stoichiometry. To confirm the functionality and versatility of the fusion protein CP-STINGΔTM, we tested a panel of NSCLC and melanoma cancer cell lines since these two cancer types can benefit from existing immunotherapy owing to high tumor mutational burden. Intriguingly, we found that CP-STINGΔTM plays distinct roles in these cell lines depending on the levels of endogenous STING expression. Specifically, codelivery of CP-STINGΔTM and cGAMP restores the STING signaling in cancer cells either naturally deficient for STING expression or genetically knocked out by CRISPR, indicating that CP-STINGΔTM and cGAMP forms a functional complex in this setting. On the contrary, CP-STINGΔTM serves as a chaperon to markedly promote the delivery of cGAMP in cells with down-regulated STING expression, requiring a 100-fold lower concentration of cGAMP than free cGAMP in STING activation and subsequent type I IFN induction. To explore the potential translation of the platform, we further confirmed potent T cell proliferation and anti-tumor immune responses *ex vivo* and extended the observation in vivo using a mouse model of vaccination. Finally, we investigated the translational potential of our platform in combination with the ICB using a syngeneic mouse melanoma model. Collectively, our CP-STINGΔTM system may provide a new paradigm of delivering STING agonists towards vaccines and cancer immunotherapy.

The most important finding of our study is that CP-STINGΔTM in complex with cGAMP can form a functional complex to activate the endogenous STING signaling in cancer cells deficient for STING. This attribute may have critical clinical implications in melanoma and lung cancers. Notably, existing STING agonism strategies have centered around developing DNA-damaging reagents to activate tumor cells to produce endogenous cGAMP, reversing the epigenetic inhibition of STING/cGAS expression, and exogenously administering STING agonists. These approaches, however, can be hampered by the fact that endogenous STING and/or cGAS are frequently silenced in tumor cells as a mechanism to evade anti-tumor immune responses(39). Specifically, the loss of tumor-intrinsic STING expression has been shown to impair tumor cell antigenicity and susceptibility to lysis by tumor-infiltrating lymphocytes through the downregulation of MHC class I expression on the surface of cancer cells. In addition to NSCLC and melanoma, decreased expression of STING in tumor cells has been correlated with poor prognosis in patients with gastric and colon cancers(26,40). Conversely, activation of tumor-intrinsic STING signaling has been found to dictate chemotherapy-induced anti-tumor cytotoxic T cell responses (e.g., olaparib) in triple-negative breast cancer(41,42).

In comparison to many existing synthetic delivery systems, our CP-STING protein as a delivery vehicle is unique in several aspects: (1) Instead of electrostatic complexation, which is particularly challenging to dinucleotides owing to low charge densities, we have made use of the inherent strong affinity between the C-terminus of STING and its agonist to efficiently encapsulate STING agonists. (2) The CP-STINGΔTM itself is in essence a single long polymer with a fixed degree of “polymerization”, and therefore is structurally well defined as evidenced by size exclusion chromatography and SDS-PAGE. This feature may minimize batch-to-batch variations, commonly occurring in synthetic delivery vehicles. (3) The fusion protein can be produced and purified from the standard *E. coli* based recombinant protein expression system in a high yield in conjunction with the low-cost metal affinity purification, which is easily accessible to many laboratories.

Future studies can involve comprehensive pharmacological characterization of the CP-STINGΔTM in the setting of systemic delivery to optimize the dose and frequency of the fusion protein. Additionally, by employing transgenic mouse models with STING deficiency in different cell types (e.g., tumor cells versus different immune cell subtypes), we can further elucidate exact targets of CP-STINGΔTM, and therefore assess the contribution of tumor-intrinsic STING in developing anti-tumor immune responses. Finally, given the modularity of the fusion protein, we can potentially substitute the cell-penetrating domain with a more specific protein domain such as nanobody to target particular cell type or TME such that our fusion platform can be extended to targeted delivery of STING agonists in a manner similar to antibody drug conjugates(43). Alternatively, direction fusion of a nanobody such as anti-PD (L)1 with STINGΔTM may simultaneously leverage ICB and STING in a single protein format. Therefore, our approach may offer a unique direction towards the STING-based therapeutics.

## MATERIALS AND METHODS

### Chemicals and antibodies

2’3’-cyclic-GMP-AMP (cGAMP) is a generous gift from Dr. Pingwei Li at Texas A&M University. Tween-20, Triton X-100, Triton X-114 were all purchased from Sigma-Aldrich (St Louis, MO). Carboxyfluorescein succinimidyl ester (CFSE) was purchased from Tonbo Biosciences (San Diego, CA). All other chemicals were purchased from ThermoFisher (Waltham, MA) and used as received. Human CXCL10/IP-10 and mouse CXCL10/IP-10 ELISA Kit, Murine TNF-alpha, and Murine IFN-gamma were respectively purchased from R&D system (Minneapolis, MN) and Peprotech (Rocky Hill, NJ). Zombie Dyes, Alexa647 anti-DYKDDDDK Tag Antibody (Clone L5), APC anti-mouse CD8a (Clone 53-6.7), FITC anti-mouse CD3 (clone 145-2C11), PerCP-Cy5.5 anti-mouse CD4 (Clone 129.29), PE anti-mouse CD8a (clone 53-6.7), PerCP-Cy5.5 cd11b (Clone M1/70), FITC anti-mouse cd11c (Clone N418), PE anti-mouse CD45 (clone 30-F11), Alexa 488 anti-mouse CD45 (clone 30-F11), FITC anti-human HLA-A,B,C Antibody (clone W6/32), FITC anti-mouse H-2Kb/H-2Db Antibody (Clone 26-8-6) were from Biolegend (San Diego, CA). Primary antibodies of STING/TM173 (D2P2F), alpha-Tubulin (DM1A), TBK1/NAK (D1B4) were from Cell signaling technology (CST, Danvers, MA). Secondary antibodies of goat anti-rabbit IgG-HRP and goat anti-mouse IgG-HRP are from Santa Cruz Biotech (Santa Cruz, CA). InVivoMAb anti-mouse PD-1 (CD279) was purchased from BioXCell (Lebanon, NH).

### Expression and purification of STINGΔTM protein variants

The human STINGΔTM protein (139-379aa) and mouse STINGΔTM (138-378aa) variants were synthesized by gblock (IDT, Coralville, IA), and cloned into pSH200 vector (a generous gift from Prof. Xiling Shen at Duke University) containing a 6xhistidine tag (His-tag), between *NcoI* and *NotI* sites. Mutants were generated with site-specific mutagenesis based on the human STINGΔTM plasmids. All plasmids were confirmed by sequencing. STINGΔTM variants were expressed as His-tag proteins from BL21 (DE3) Escherichia coli (*E. coli*). All proteins were expressed as cultures grown in Luria-Bertani broth (LB) (5g sodium chloride, 5g tryptone, 2.5g yeast extract, and 500 mL of distilled water), supplemented with 100 μg/ml Ampicillin. After outgrowth at 37°C with 225 rpm in a shaker, and until optical density (OD600) reached 0.6, 1 mM IPTG was added to induce the protein expression for 16 to 18 hours at 20°C and 225 rpm. Cells were then collected by centrifugation at 5000x g for 20 minutes at room temperature. The bacterial pellets were resuspended in a 10 mL protein binding buffer (50 mM sodium phosphate, 0.5 M sodium chloride, 10 mM imidazole) and stored at −80°C until purification. The frozen cultures were thawed and lysed with 1% Triton-100, 1 mg/mL lysozyme, 1mM PMSF, and one EDTA-free protease inhibitor cocktail tablet at room temperature for 20 min. The lysate was disrupted by ultrasonication at 5-second intervals for a total of 5 min each at 18 W on ice. Insoluble debris was removed by centrifugation at 12000x g for 60 min, at 4 °C. Protein purification was carried out by affinity chromatography using Cobalt agarose beads. 10 mL of raw protein extracts were applied to the protein binding buffer-equilibrated beads, followed by three washes with protein binding buffer plus 0.1% Triton-114 for endotoxin removal. After elution (50 mM sodium phosphate, 0.5 M Sodium chloride, 150 mM imidazole), protein extracts were loaded to fast protein liquid chromatography (FPLC, NGC Quest 10 Chromatography System, Biorad) for 3X PBS buffer exchange and purification. Protein fractions detected at λ = 280 nm were collected. Purified STINGΔTM variants concentrations were determined by DC protein assay and purities were verified by SDS-PAGE. Protein aliquots were kept at −80°C at all times until further use.

### Animal work

All work with C57BL/6J mice (females, 7-10 weeks old) and OT-1 transgenic mice (The Jackson Laboratory, Bar Harbour, ME) was performed in accordance with institutional guidelines under protocols of NU-20-0312R (C57BL/6J) and NU-19-0106R (OT-1) approved by Northeastern University-Institutional Animal Care and Use Committee (NU-IACUC). All mice were maintained in a pathogen-free facility following the National Research Council of the National Academies.

### Cell lines and cell culture

Non-small cell lung cancer cell lines A549, H1944, and H2122 harboring KRAS/LKB1 comutations and H1944 Knockouts (H1944 STING-knockout, H1944 cGAS-knockout, H1944 scramble-knockout) were generous gifts from Dr. David Barbie’s lab. Human and murine cell lines of B16F10, HeLa, HEK293T, SK-MEL-3, and SK-MEL-5, were obtained from the American Type Culture Collection (ATCC, Rockville, MD). Yummer1.7 was requested from the Koch Institute (Cambridge, MA). B16-OVA(257-264aa) and Yummer1.7-OVA(257-264aa) were generated through transfection with plasmids encoding full lengths of OVA and EGFP, and sorted by FACS for GFP expression. A549, SK-MEL-3, SK-MEL-5, Yummer1.7, HeLa, and HEK293T were cultured in Dulbecco’s modified Eagle’s medium (DMEM) supplemented with 10% fetal bovine serum (FBS), 100 U/mL penicillin-streptomycin, and 100x Non-Essential Amino Acid (NEAA). H1944, H2122, HCC44, and H23 were cultured in RPMI-1640 supplemented with 10% FBS, 100 U/mL penicillin-streptomycin, and 100x NEAA. H1944 STING-knockout, H1944 cGAS-knockout, and H1944 scramble-knockout were cultured in RPMI-1640, with 10% FBS, 100 U/mL penicillin-streptomycin, 100x NEAA with 1μg/mL puromycin selection. Cells were kept in a humidified incubator with 5% carbon dioxide (CO_2_) at 37°C and routinely tested mycoplasma negative by PCR. All the cell experiments were performed between passages 2 and 10.

### Lentivirus production and cell line generation

Lentiviral vector plasmids of *pFUW Ubc OVA (252-271aa) EGFP, EGFP Luciferase puro (663)* were used to generate lentiviral particles. 7.5 μg of packaging plasmid psPAX2, 2.5 μg of envelope plasmid pMD2.G, 10μg of Lentiviral vector plasmids, and 10 μL TransIT-X2 were mixed in 1 mL Opti-MEM. After 30 minutes of incubation at room temperature, the plasmid mixture was added to 70% confluency HEK293T cells. Supernatants were collected at 48 hours and 72 hours after transfection and centrifuged at 1000 g for 10 minutes to remove the debris. Harvested Lenti-viral supernatants were kept at −80 C until further cell line generation. After targeted cell lines of B16F10 and Yummer 1.7 reached 70% confluency, lentiviral supernatants were added to the cells with 8 μg/mL polybrene. Transfected cells were selected with 1 μg/mL puromycin.

### Enzyme-linked immunosorbent assay (ELISA)

For human CXCL10 and mouse CXCL10, cells (1 to 2×10^4^) were cultured with premixed complexes of 40 μg/mL, or 10 μg/mL STINGΔTM variants with or without 1 μg/mL or 0.25 μg/mL cGAMP for 72 hours. Conditioned supernatants were collected for ELISA quantification according to the manufacturer’s instructions. Values represent the average of four to six replicates from at least two independent experiments. For analysis of anti-OVA IgG level, we conducted the ELISA as previously described(44). For cytokine quantification in the treatment study, tumors were harvested and grounded in tissue protein extraction reagent (T-PER^TM^) with 1% proteinase inhibitors. The lysates were incubated at 4 °C for 30 min with rotation. The supernatant from each lysate was collected after removing debris through centrifugation. The quantifications of CXCL10, TNF-α, and IFN-γ were performed according to the manufacturer’s instructions.

### Immunofluorescence staining

A549, H1944, and HeLa were seeded in chamber slides at a density of ~5×10^4^ 24 hours before incubation with 40 μg/mL STINGΔTM variants and 1 μg/mL cGAMP complexes. After another 24 hours, cells were washed with PBS once, and fixed with 70% ethanol. After permeabilization with PBS containing 0.1% Triton X-100 for 15 minutes, cells were washed and incubated with the anti-DYKDDDDK Tag antibody at 1:500 dilution in 1xPBS with 1% BSA and 0.05% TWEEN 20 (PBST) at 4°C overnight. Cells were then washed for 30 minutes in PBST, and incubated with Alexa488-Phalloidin (CST) in 1:100 dilution for 1 hour. After washing cells with PBST for three times for 10 minutes each, cells were counter-stained with DAPI in mounting media at room temperature. Images of the cells were visualized and captured by Nikon Eclipse Ti2 microscope (Tokyo, Japan) and analyzed by ImageJ (NIH).

### Fluorescence imaging analysis

Three days after injection with complexes, tumors were harvested and placed in OCT in tissue cassettes and frozen on ice for cutting into 8-10μm sections in slides. The slides were washed with PBS for 10 min at room temperature, dried on a paper towel and incubated with anti-CD45 diluted in the antibody buffer (10% FBS in PBS) for 1 hour at room temperature in the dark. After three washes with PBS, the slides were fixed in 4% paraformaldehyde in PBS. Slides were incubated with 0.025% saponin in PBS for permeabilization. Anti-DYKDDDDK were added on the sections for overnight incubation at 4 °C in the dark. Slides were washed in PBS with 0.0025% saponin for 10 min twice. After incubating with secondary antibody for 1 hour in the dark, slides were rinsed with PBS with 0.0025% saponin and counterstained with DAPI. The stained tumor slides were imaged using a Nikon microscope.

### Flow cytometry

For uptake study, 1×10^5^ cells were seeded in 12-well plates in their corresponding complete culture medium and incubated for 24 hours. After treatment with 40 μg/mL STINGΔTM variants with or without 1 μg/mL cGAMP for 24 hours, cells were washed with PBS and treated with trypsin for at least 15 minutes to remove STING proteins nonspecifically bound to the cell surface. Cells were transferred to 96-well v-bottom plates and collected through 300xg centrifugation for 3 minutes. After twice washes with 200 μl PBS, cells were fixed with 70% ethanol for 20 minutes. The fixed cells were washed with PBS for 10 minutes three times. Cells were resuspended in anti-DYKDDDDK Tag Antibody at 1:1000 dilution in antibody dilution buffer (1xPBS containing 1% BSA and 0.05% Tween 20) and incubated for 2 hours at room temperature in the dark. Antibodies were removed by rinsing cells with PBST three times. The cell suspension in PBS was loaded to Attune flow cytometry (Thermofisher, Waltham, MA). Doublets and dead cells were excluded before analysis.

For *in vitro* MHC-I analysis, 10000 cells were incubated with 40 μg/mL STINGΔTM variants and 1 μg/mL cGAMP in a complete culture medium for 48 hours before staining. Cells were rinsed by PBS, detached by 100μl 5mM EDTA in PBS with a fixable live/dead dye, NIR Zombie Dye (Biolegend), at 1:1000 dilution for dead cell exclusion. After staining was quenched by FACS buffer (5% FBS, 2mM EDTA, 0.1% sodium azide in PBS), cells were resuspended by FACS buffer containing 0.4μg/ml anti-human HLA-A,B,C antibody or FITC anti-mouse H-2Kb/H-2Db antibody, and incubated on ice for 30 min in the dark. Stained cells were washed twice and resuspended in the FACS buffer for flow cytometric analysis in FlowJo (Franklin Lakes, NJ). After excluding doublets and debris of dead cells, gating strategies determined through control staining were applied for analysis while compared with FITC Mouse IgG2a, κ Isotype control antibody stained cells.

For OT-1 CD8+ T cells stimulation, CFSE stained lymphocytes were collected through 500x g centrifuge for 3 min and washed with 200μl PBS. 100μl Zombie dye in PBS at 1:1000 dilution was added to the lymphocyte and incubated for 30 min at room temperature avoiding light. Zombie dye staining was quenched by 100μl FACS buffer. After 3 min centrifuge at 500x g, OT-1 CD8^+^ T cells were selected by 100μl APC antimouse CD8a Antibody in FACS buffer at 1:1000 dilution after 30 min incubation on ice. Co-stained cells were resuspended in the FACS buffer and quantified under the flow cytometer.

For *in vivo* tumor profiling, dissected tumors were digested in 1 mg/mL collagenase D for 1 hour at 37 °C. Single-cell suspensions were obtained from mincing the tumor through a 70 μm cell strainer. After staining with NIR zombie dye for dead cell exclusion, cells were neutralized and blocked with anti-CD16/CD32 for 5 minutes on ice and stained with antibodies against surface markers CD45, CD3, CD4, CD8, CD11b, CD11c on ice for 30 minutes in FACS buffer. For intracellular staining, cells were fixed, permeabilized, and stained with anti-DYKDDDDK tag antibody. All samples were analyzed by FlowJo after loading to the flow cytometer.

### Cell viability assay

The effects of STINGΔTM variants and cGAMP complexes on cell viability were determined by 3-(4,5-dimethylthiazol-2-yl)-2,5-diphenyltetrazolium bromide (MTT) assay. 1000 cells were seeded in 96-well plates and treated with 40μg/mL STINGΔTM variants and 1μg/mL cGAMP for 120 hours in 5% CO2 at 37°C in a humidified incubator. Cells were further incubated with 0.5 mg/mL MTT dissolved in sterilized 1x PBS at 37°C for 2 hours before DMSO was added into each well to dissolve formazan crystals. The absorbance of each well was determined at 570 nm on an automated Bio-Rad microplate reader (Bio-Rad Laboratories, Hercules, CA). Untreated cells as control were considered to be 100% viable.

### Lymphocyte preparation from lymph nodes in OT-1 mice

The mesenteric, inguinal, axillary, and brachial lymph nodes dissected from OT-1 mouse were homogenized to generate a single cell suspension, and the released cells in lymphocyte growth medium (RPMI1640 complete media and 50μM 2-mercaptoethanol) were pelleted and resuspended in 10ml PBS. The lymphocyte was washed and stained with 1μM CFSE in 1x PBS for 20 min until the staining was terminated by 10% FBS. The stained lymphocyte was resuspended and cultured in lymphocyte growth medium in a humidified incubator to release excessive CFSE. After 2 hours incubation, lymphocyte was collected and resuspended in lymphocyte growth medium with 20U/ml interleukin (IL)-2.

### Coculture of OT1 lymphocytes with B16-OVA or YUMMER 1.7-OVA

100μl of 1x 10^6^ lymphocytes in lymphocyte growth medium with 20U/ml IL-2 was added into the 96-well plate with 100μl of 1×10^4^ B16-OVA(257-264aa) treated with STINGΔTM variants with or without cGAMP 48 hours ahead. On days 3, 100μl of lymphocytes were gently collected for flow cytometry analysis. 100μl fresh lymphocyte growth medium with 20U/ml IL-2 was added to each well for leftover lymphocyte growth. On day 5, after lymphocytes were collected, B16-OVA(257-264aa) attached wells were washed with PBS twice for subsequent MTT assay.

### Immunizations, tumor inoculation and treatment in mice

Analysis of immunizations for adjuvant potential performed in C56BL/6 mice with B16-OVA (257-264aa) was conducted as previously described(44). For treatment study, one million Yummer1.7 cells in 100μl Opti-MEM were subcutaneously injected into the flank of mice. At 6-9 days later, when tumors reached 150 mm^3^ in volume, animals were injected intratumorally with ~25ul vehicle control, 2.5 μg cGAMP only or 100 μg STINGΔTM variants and 2.5 μg cGAMP complex in Opti-MEM.

### Statistical analysis

Statistical significance was evaluated using one-way ANOVA followed by Tukey *post hoc* test. P values less than 0.05 were considered significant. Statistical significance is indicated in all figures according to the following scale: *P<0.05, **P<0.01, ***P<0.001, and ****P<0.0001. All graphs are expressed as the means ± SEM. In one-way ANOVA followed by *post hoc* tests, we marked asterisks only in pairs of our interest.

## ACKNOWLEDGEMENTS

This work was supported by the Northeastern University Faculty start-up funding (JL), Northeastern University-Dana Farber Cancer Institute Cancer Drug Development (JL and DAB), and Peer Reviewed Medical Research Program from the Department of Defense’s Congressionally Directed Medical Research Programs (JL). We are grateful to Dr. Yingzhong Li at the Koch Institute of Integrative Cancer Research at MIT for scientific advice. We would like to express our gratitude to Professor Sara Rouhanifard at Northeastern University’s Department of Bioengineering for generously sharing her lab’s fluorescent microscope; Professor Ke Zhang at Northeastern University’s Department of Chemistry for sharing his lab’s equipment.

