## supplemental information for "Engineering the Immune Adaptor Protein STING as a Biologic" (from another version of this preprint: version 1, 2021-02-18)

### Supplementary Information

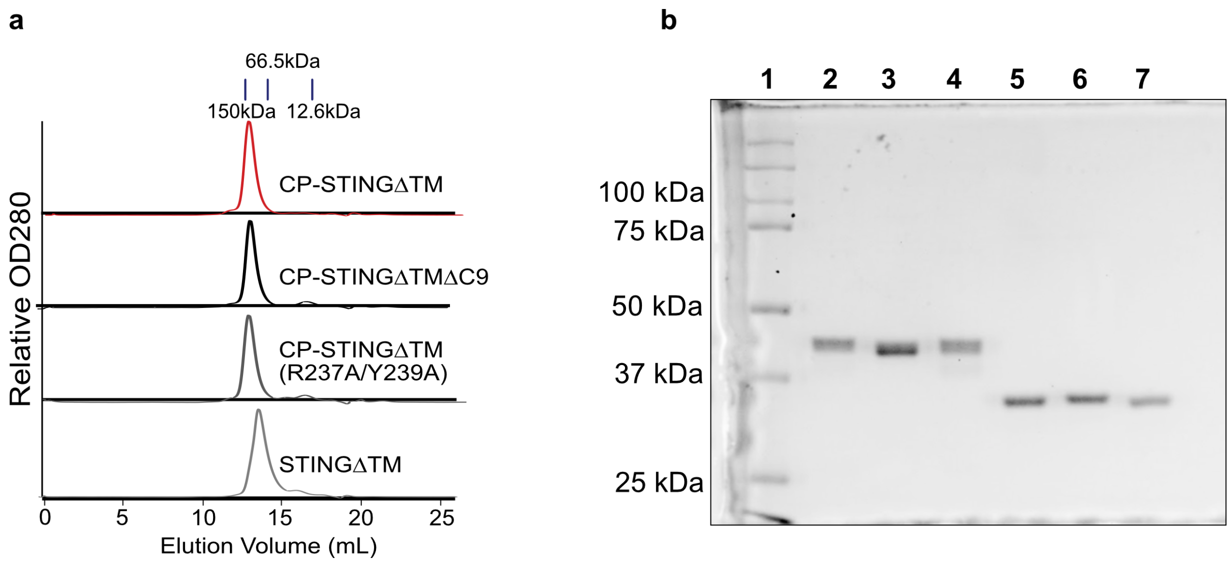


**Figure S1**. (**a**) Size exclusion chromatography (SEC) of CP-STINGΔTM, CP-STINGΔTMΔC9, CP-STINGΔTM (R237A/Y239A) and STINGΔTM in PBS buffer. (**b**) SDS-PAGE of CP-STINGΔTM (Lane **2**), CP-STINGΔTMΔC9 (Lane **3**), CP-STINGΔTM (R237A/Y240A) (Lane **4**), STINGΔTM (Lane **5**), STINGΔTMΔC9 (Lane **6**), and STINGΔTM(R237A/Y239A) (Lane **7**) under a denaturing condition.

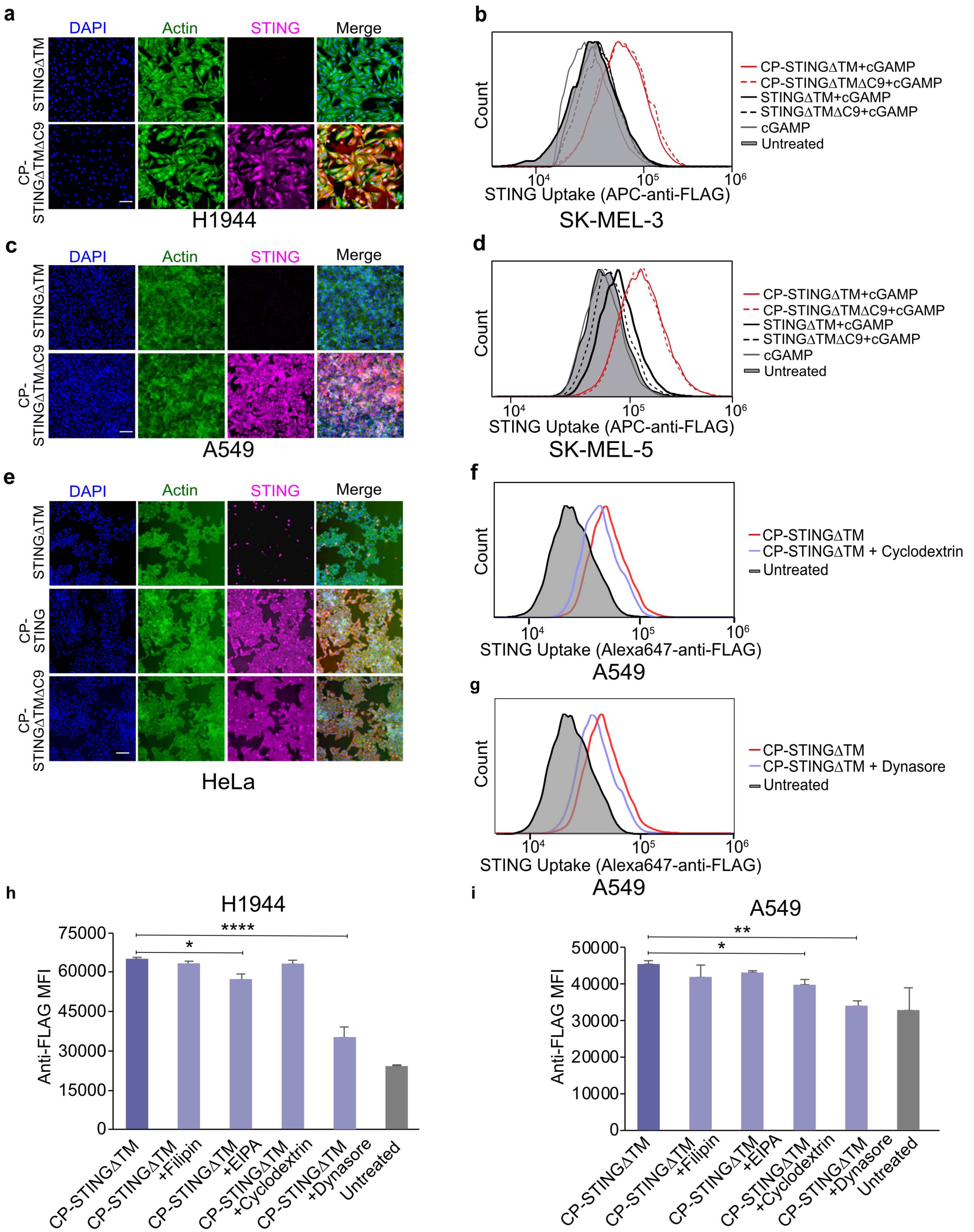


**Figure S2**. **CP-STINGΔTM effectively internalizes cancer cells**. Fluorescence microscopy imaging of internalized CP-STINGΔTMΔC9 in H1944 ($STING_{low}$) **(a)**, A549

(STING$_{absent}$) **(c)** and ovarian cell line HeLa (**e**) (scale bar = 100 μm). Flow cytometry of internalized CP-STINGΔTM in SK-MEL3 (STING$_{positive}$) (**b**) and SK-MEL5 (STING$_{absent}$) (**d**). Effects of indicated small molecule inhibitors on the cellular uptake were performed in A549(STING$_{absent}$) (**f and g**). Alexa647-Anti-FLAG MFI of cellular uptake inhibition by small molecule inhibitors in (**h)** H1944 (STING$_{low}$) and (**i**) A549 (STING$_{absent}$) under flow cytometry analysis.

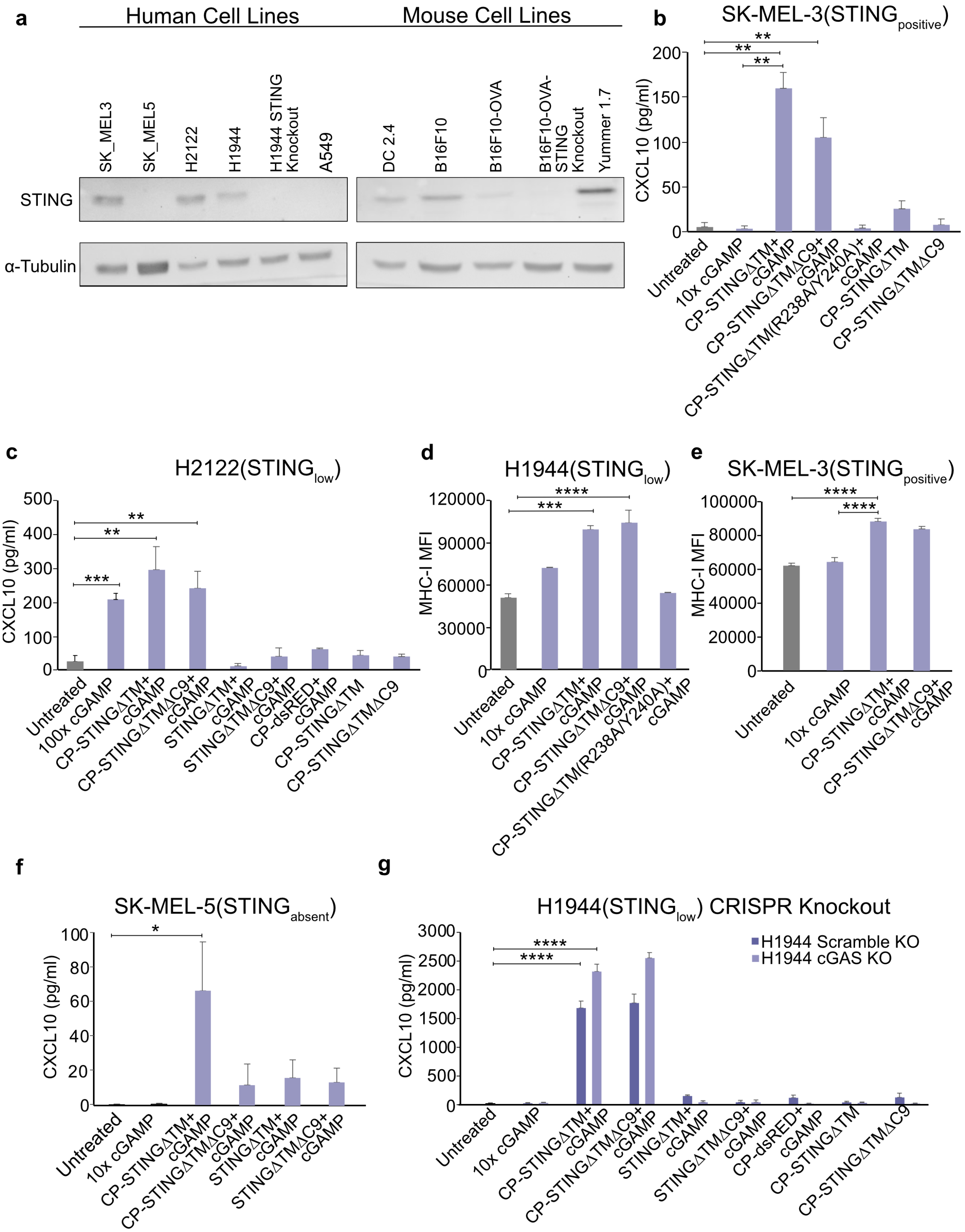


**Figure S3**: (**a**) Immunoblotting of endogenous STING in human and mouse cell lines. CP-STINGΔTM plays a chaperon role in enhancing cGAMP delivery and subsequent CXCL-

10 production in (**b**) SK-MEL-3 ($STING_{positive}$) and (**c**) H2122 ($STING_{low}$), while CP-STINGΔTM + cGAMP forms a functional complex in (**f**) SK-MEL-5 ($STING_{absent}$). Similarly, quantification of MHC-I upregulation in different combinations in (**d**) H1944 ($STING_{low}$) and (**e**) SK-MEL-3 ($STING_{positive}$) indicates the chaperon role of CP-STINGΔTM. (**g**) The endogenous cGAMP is not required for enhanced delivery of exogenously administered "CP-STINGΔTM + cGAMP" in H1944 cells, in which the cGAS was knocked out by CRISPR.

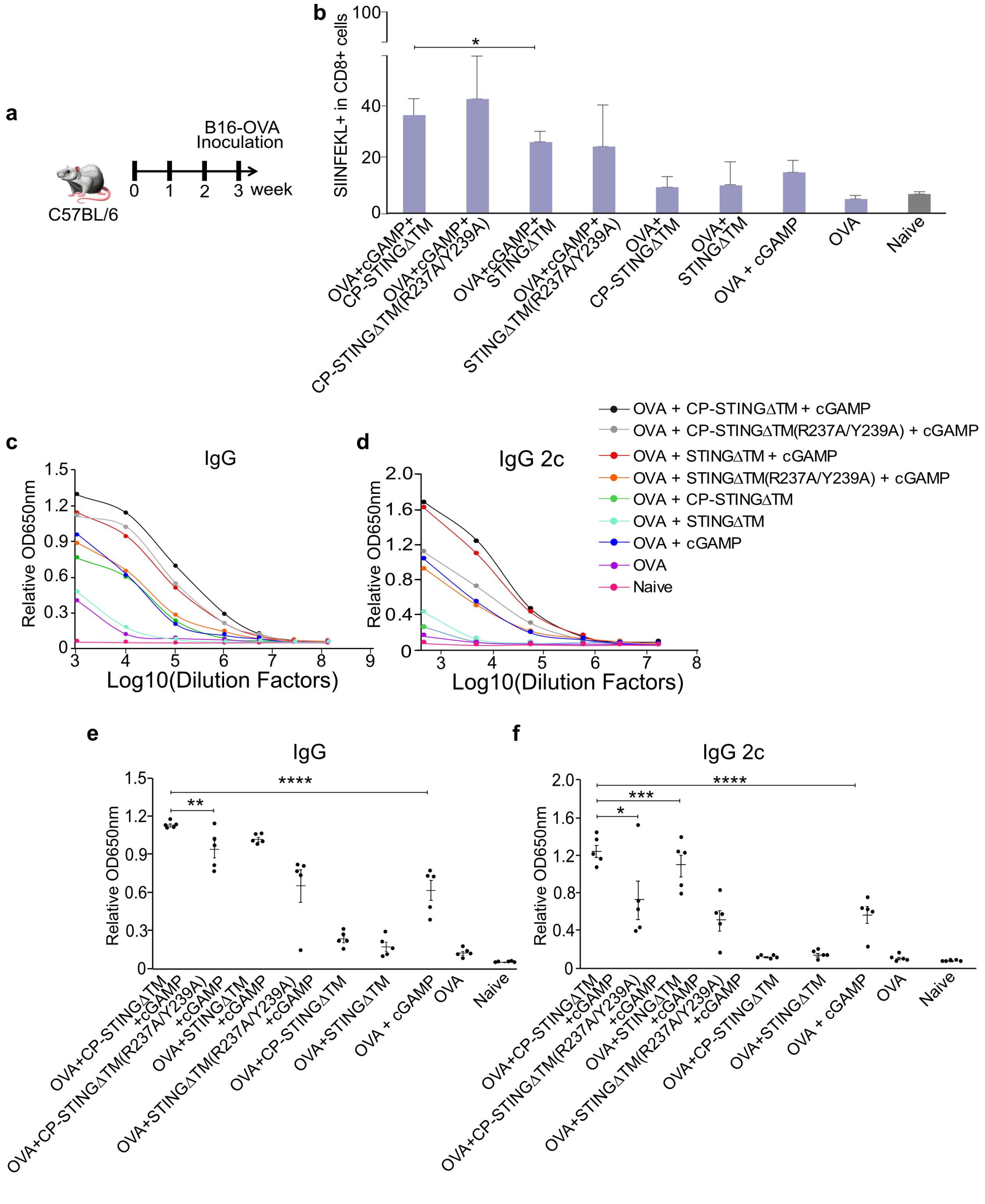


**Figure S4**. (**a**). Vaccination strategy in this study. (**b**) Percentage of CD8 T cells carrying the MHC-I-SIINFEKL epitope from $OVA_{257\text{-}264aa}$ via tetramer staining. (**c** and **d**) OVA-specific IgG and IgG 2c antibody levels in mouse serum in different treatment groups were measured by ELISA. (**e** and **f**) Representative plots of OVA-specific IgG and IgG2c in serum from each mouse in different treatment groups.

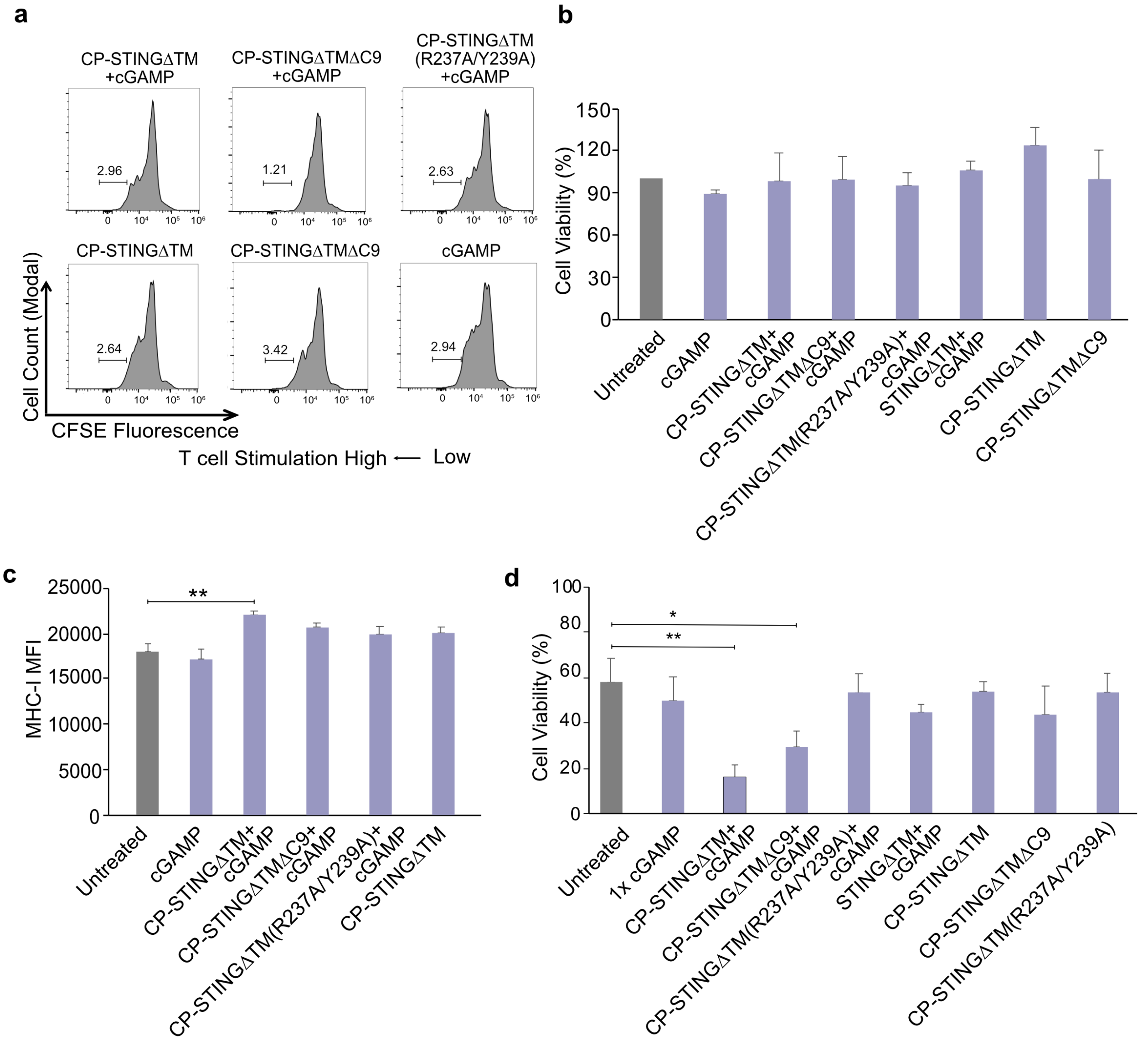


**Figure S5**. (**a**) T cell stimulation and (**b**) B16-GFP tumor cell viability after coculturing with OT1 cells for 120 hours. (**c**) MHC-I upregulation in B16-OVA (257-264aa) by flow cytometry. (**d**) Yummer1.7-OVA (257-264aa) cell viability after coculturing with OT1 cells for 120 hours.

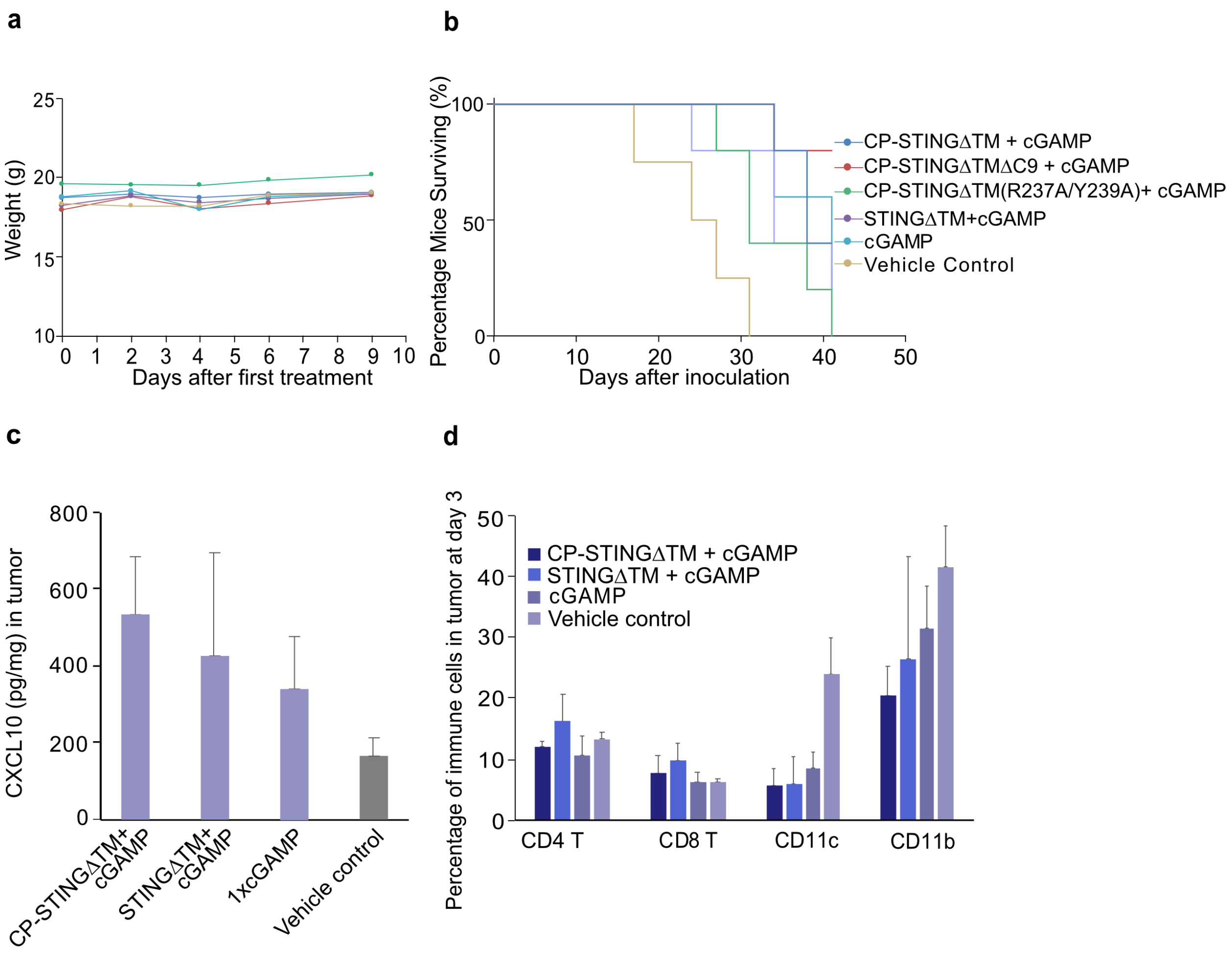


**Figure S6**. (**a**) Body weight measurement in different treatment group over the course of treatment. (**b**) Survival curves of mice. (**c**) CXCL10 expression induced by different treatment groups (n=3) was quantified by ELISA. (**d**) Immune cell profiling via antibody staining for CD4, CD8, CD11c and CD11b in YUMMER 1.7 tumors receiving different treatment regimens.

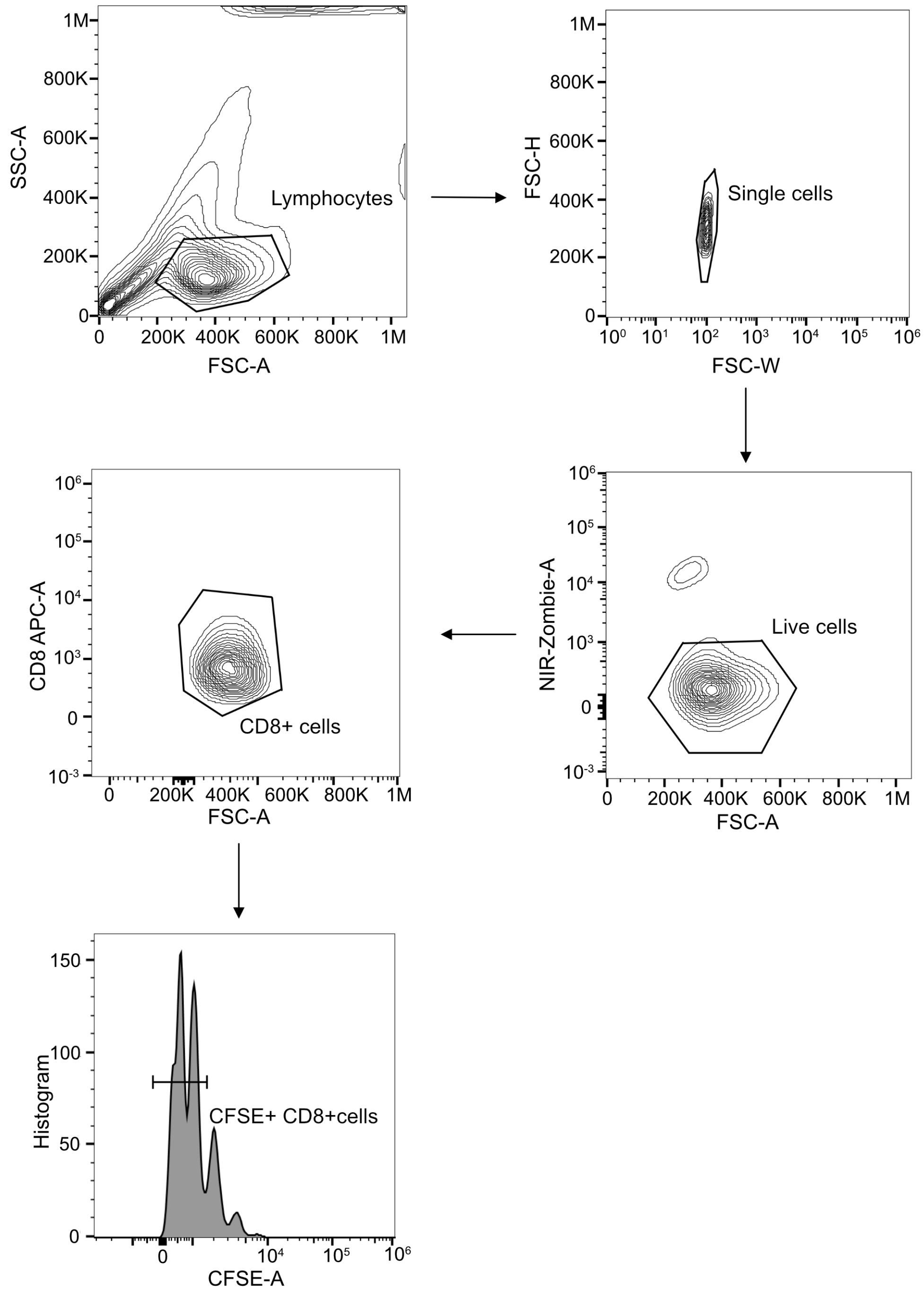


**Figure S7**. Gating strategies of analyzing OT1 T cell proliferation in the *ex vivo* model
